# Faecal DNA to the rescue: Shotgun sequencing of non-invasive samples reveals two subspecies of Southeast Asian primates to be Critically Endangered species

**DOI:** 10.1101/867986

**Authors:** Andie Ang, Dewi Imelda Roesma, Vincent Nijman, Rudolf Meier, Amrita Srivathsan, Rizaldi

**Author notes:** Co-first (Andie Ang, Dewi Imelda Roesma).

## Abstract

A significant number of Southeast Asian mammal species described in the 19^th^ and 20^th^ century were subsequently synonymized and are now considered subspecies. Many are affected by rapid habitat loss and there is thus an urgent need to re-assess the conservation status based on species boundaries established with molecular data. However, such data are lacking for many populations and subspecies. We document via a literature survey and empirical study how shotgun sequencing of faecal DNA is a still underutilized but powerful tool for accelerating such evaluations. We obtain 11 mitochondrial genomes for three subspecies in the langur genus *Presbytis* through shotgun sequencing of faecal DNA (*P. femoralis femoralis*, *P. f. percura*, *P. siamensis* cf. *cana*). The genomes support the resurrection of all three subspecies to species based on multiple species delimitation algorithms (PTP, ABGD, Objective Clustering) applied to a dataset covering 40 species and 43 subspecies of Asian colobines. For two of the newly recognized species (*P. femoralis*, *P. percura*), the results lead to an immediate change in the IUCN status to Critically Endangered due to small population estimates and fragmented habitat. We conclude that faecal DNA should be more widely used for clarifying species boundaries in endangered mammals.

## Introduction

Human impacts on the environment have rapidly accelerated species extinction via habitat degradation and climate change. Recent report by Intergovernmental Science-Policy Platform on Biodiversity and Ecosystem Services (IPBES) predicts that climate change has already affected the distribution of nearly half (47%) of land-mammals^1^. Conservation efforts are urgently needed but are hampered by the lack of data for a large number of mammal species, subspecies, and populations which face imminent extinction^2,3,4^. A typical example is Asian primates for which 70% of the species are threatened with extinction^5^. Effective conservation programs are needed but they require a robust understanding of species numbers and boundaries based on up-to-date taxonomic information^6,7^. Unfortunately, this information is lacking for many rare, globally threatened, and elusive mammalian species. Many lack molecular data and collecting these data is difficult because invasive sampling that would yield fresh tissues is often not feasible.

This leaves only three alternative sources of DNA. The first is museum specimens, but the number of samples in museums tends to be small and many were collected in the 19^th^ or early 20^th^ century thus reflecting (historic) genetic diversity prior to extensive habitat loss. The second is tissue samples obtained from specimens that died of “natural causes” such as road accidents. The third source of genetic material is non-invasive samples such as hair and faeces. Arguably, faecal samples are still an underappreciated source of information although they could be collected in good numbers during routine field surveys. This can make faecal samples particularly useful for data-deficient taxa that are in urgent need for re-assessment of species boundaries. Faecal samples contain a complex pool of DNA including that of the host. The host DNA is particularly informative because it reflects the current genetic diversity of the species. However, many field research protocols still lack the collection of faecal samples although it is now straightforward to obtain complete mitochondrial genomes from such samples using shotgun sequencing^8,9^. In this study, we document the power of faecal metagenomics for testing the species boundaries in two species of Asian primates that are listed as Data Deficient on the IUCN Red List of Threatened Species.

Asian colobines (langurs and odd-nosed monkeys) are a diverse group of mammals, with 55 recognized species (87 spp.) belonging to seven genera (*Nasalis*, *Presbytis*, *Pygathrix*, *Rhinopithecus*, *Semnopithecus*, *Simias*, *Trachypithecus*)^10^; i.e., nearly half of all primate species in Asia are colobines. Unfortunately, many of these species are dependent on habitats that are quickly disappearing. Thus, nine species are already considered Critically Endangered, 23 are Endangered, and nine Vulnerable according to IUCN threat criteria^5^. This also applies to the genus *Presbytis* which is one of the most species-rich primate genera^11^. The 17 recognized species are found in the tropical rainforests of Sundaland, including the Malay Peninsula and the western Indo-Malay Archipelago^12,13^. Eleven species within *Presbytis* (*chrysomelas*, *comata*, *femoralis*, *frontata*, *hosei*, *melalophos*, *natunae*, *potenziani*, *rubicunda*, *siamensis*, and *thomasi*) were recognized in the last IUCN assessment^14^. The assessment predated the elevation of six subspecies to species (see Roos et al.^10^: *bicolor*, *canicrus*, *mitrata*, *sabana*, *siberu*, *sumatrana*) which suggests that many of the *Presbytis* taxa currently ranked as subspecies are in urgent need for re-assessment with molecular data. Unfortunately, these data are lacking for many subspecies and species which has serious consequences for the proper conservation assessment of these taxa.

Meyer et al.^11^ presented the most comprehensive phylogenetic reconstruction of *Presbytis* which included 13 of the 17 recognized species. The analysis was based on two mitochondrial markers (cyt-b and d-*loop*). Of interest in this study is the banded langur *P. femoralis*, the relationship within which has been addressed by several studies*. Presbytis femoralis* is found on the Malay Peninsula and the island of Sumatra^15^ (Fig. 1). The species currently consists of three subspecies that were originally described as species because they are distinguishable based on a combination of morphological characters. However, many primatologists currently considered these characters insufficient for recognizing the three taxa as species (see Table 1).

**Table 1.**
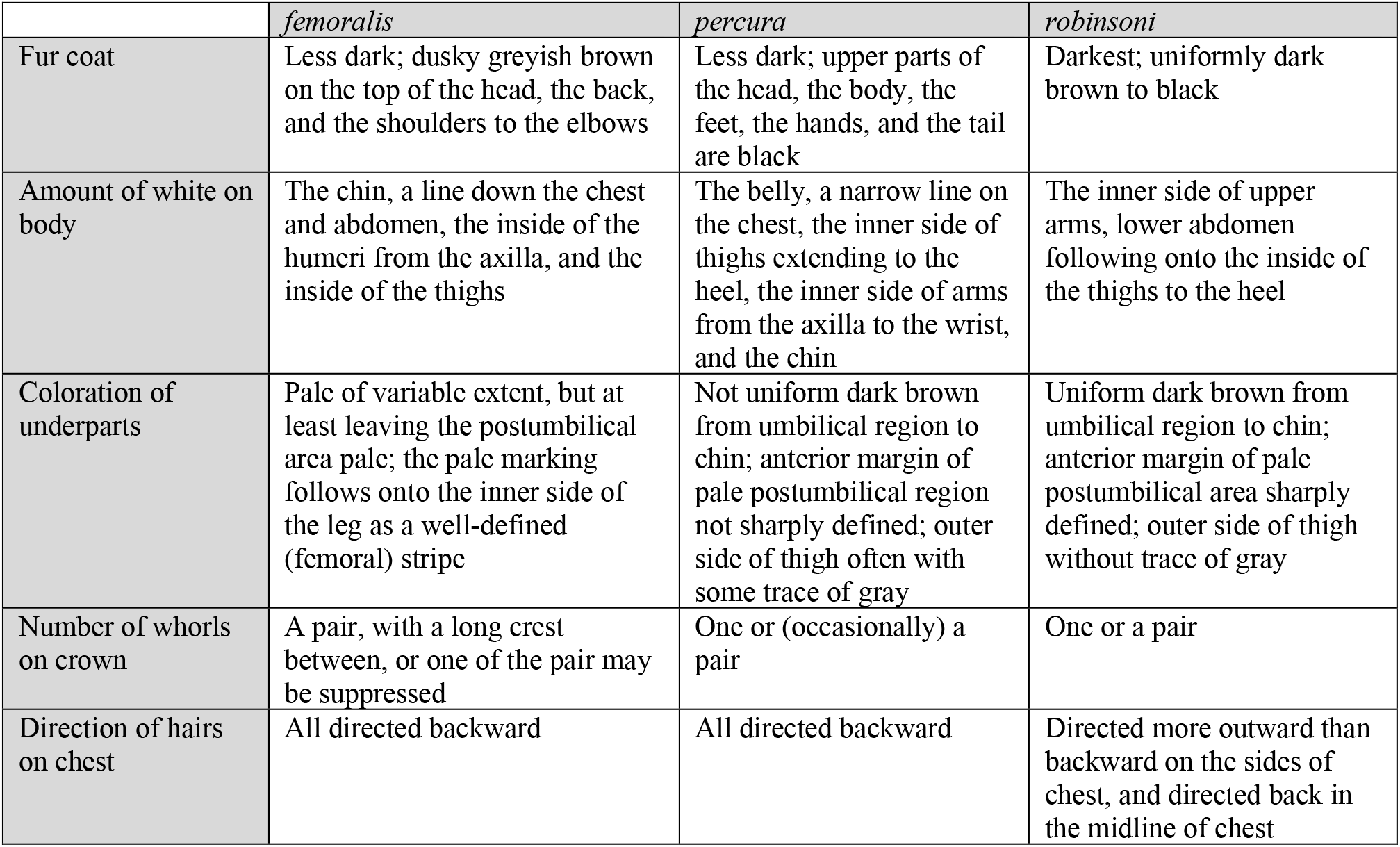
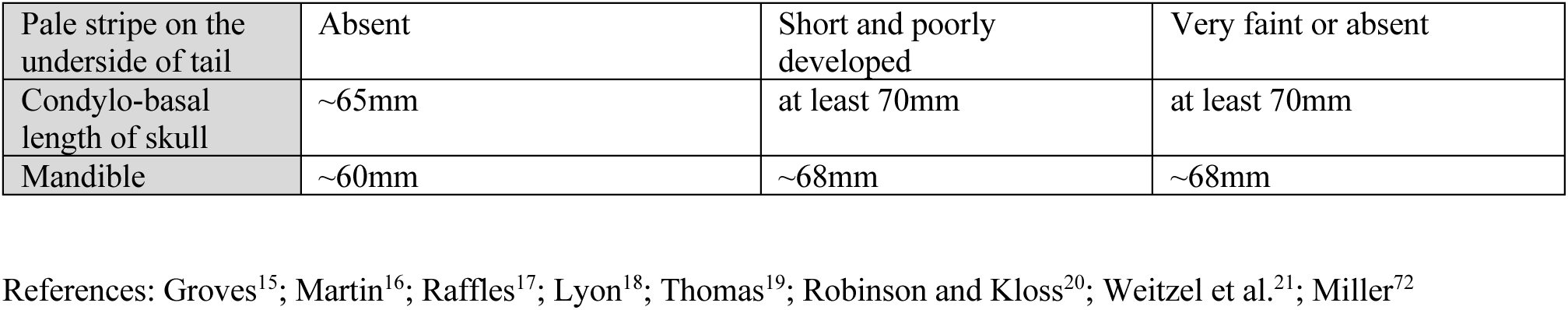
Morphological characters of three subspecies of *Presbytis femoralis*

|  | *femoralis* | *percura* | *robinsoni* |
| --- | --- | --- | --- |
| Fur coat | Less dark; dusky greyish brown on the top of the head, the back, and the shoulders to the elbows | Less dark; upper parts of the head, the body, the feet, the hands, and the tail are black | Darkest; uniformly dark brown to black |
| Amount of white on body | The chin, a line down the chest and abdomen, the inside of the humeri from the axilla, and the inside of the thighs | The belly, a narrow line on the chest, the inner side of thighs extending to the heel, the inner side of arms from the axilla to the wrist, and the chin | The inner side of upper arms, lower abdomen following onto the inside of the thighs to the heel |
| Coloration of underparts | Pale of variable extent, but at least leaving the postumbilical area pale; the pale marking follows onto the inner side of the leg as a well-defined (femoral) stripe | Not uniform dark brown from umbilical region to chin; anterior margin of pale postumbilical region not sharply defined; outer side of thigh often with some trace of gray | Uniform dark brown from umbilical region to chin; anterior margin of pale postumbilical area sharply defined; outer side of thigh without trace of gray |
| Number of whorls on crown | A pair, with a long crest between, or one of the pair may be suppressed | One or (occasionally) a pair | One or a pair |
| Direction of hairs on chest | All directed backward | All directed backward | Directed more outward than backward on the sides of chest, and directed back in the midline of chest |
| Pale stripe on the underside of tail | Absent | Short and poorly developed | Very faint or absent |
| Condyllo-basal length of skull | ~65mm | at least 70mm | at least 70mm |
| Mandible | ~60mm | ~68mm | ~68mm |

References: Groves<sup>15</sup>; Martin<sup>16</sup>; Raffles<sup>17</sup>; Lyon<sup>18</sup>; Thomas<sup>19</sup>; Robinson and Kloss<sup>20</sup>; Weitzel et al.<sup>21</sup>; Miller<sup>72</sup>

**Figure 1.**
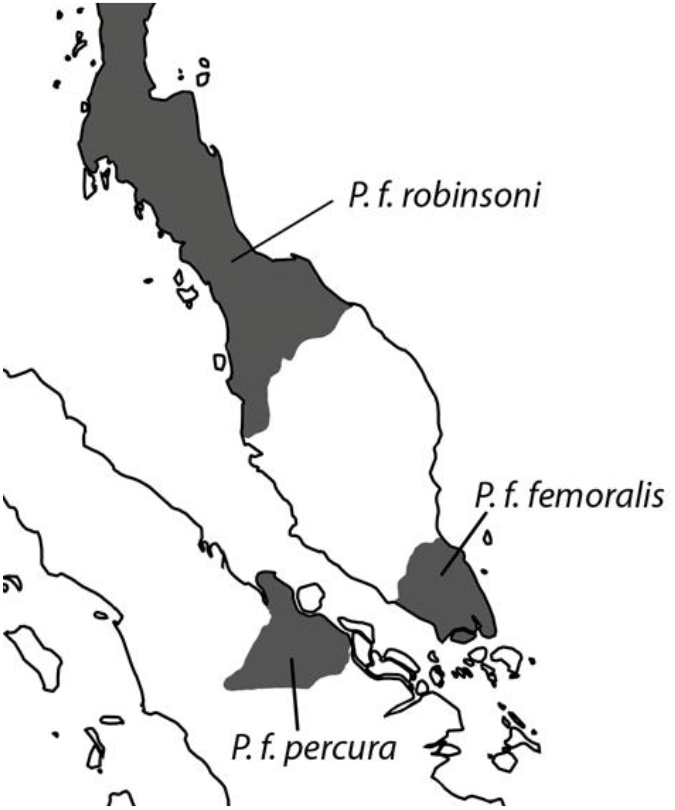
Distribution of the banded langur *Presbytis femoralis* on the Malay Peninsula and eastern Sumatra. Map: Ang Yuchen

The nominal species, *Presbytis femoralis*, was described by Martin (1838) based on specimens collected by Raffles (1821) from Singapore^16,17^. Raffles’ banded langur *P. f. femoralis* occurs in southern Peninsular Malaysia and Singapore. East Sumatran banded langur *P. f. percura* (Fig. 2) occurs only in eastern Sumatra and was described by Lyon (1908) based on specimens collected from near Siak Kecil River, Makapan, Kompei, Pulau Rupat, and Salat Rupat^18^. Robinson’s banded langur *P. f. robinsoni* was described by Thomas (1910) based on white phenotypic variants collected in Trang, southern Thailand^19,20,21^. However, typical *robinsoni* specimens are uniformly dark brown to black, with the inner side of upper arms, lower abdomen following onto the inside of the thighs to the heel being white. *Presbytis f. robinsoni* is widespread and ranges from northern Peninsular Malaysia to southern Thailand and Myanmar.

**Figure 2.**
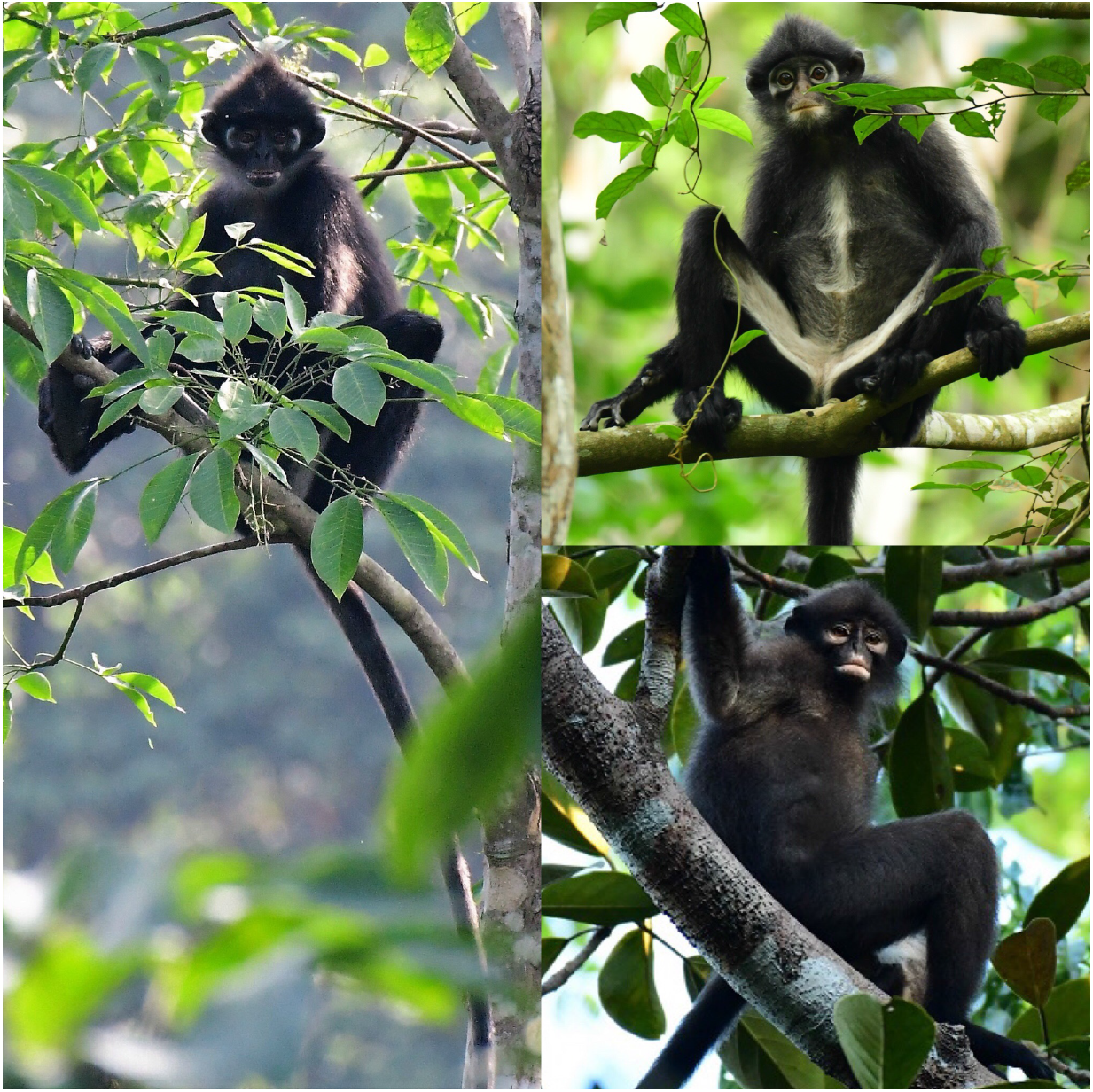
Three subspecies of *Presbytis femoralis*; clockwise from East Sumatran banded langur *P. f. percura* (1), Raffles’ banded langur *P. f. femoralis* (2), to Robinson’s banded langur *P. f. robinsoni* (3). Photos: Andie Ang

*Presbytis femoralis*, encompassing all three subspecies, is listed as Vulnerable in the most recent IUCN Red List assessment, with the nominate subspecies considered Endangered because the known populations are restricted to small and isolated patches of forest. In addition, one population from Singapore showed low genetic variability^9,22^. *Presbytis f. robinsoni* is considered Near Threatened, while the least known and least studied subspecies is *P. f. percura* which is currently Data Deficient^23^. Genetic data suggest that at least *P. f. femoralis* and *P. f. robinsoni* are different species^24,25^ which is also in agreement with the aforementioned morphological characters. However, resolving all species-level boundaries within banded langurs required data for *P. f. percura*.

For several reasons the species boundaries within *P. femoralis* remain poorly understood. The main problem is the lack of molecular data for *P. f. percura.* However, even if the data were available, a comprehensive analysis would still be difficult because the sequence data from three published analyses were not submitted to public sequence repositories (cyt-b, 12S rDNA, and d-*loop*)^26,27,28^. This means that currently the only publicly available molecular data are for *P. f. femoralis* from the type locality in Singapore (KU899140)^9^ and *P. f. robinsoni* from Redang Panjang, Malaysia (DQ355299)^29^. Some additional molecular data can be reconstructed based on a table published in Abdul-Latiff et al.^28^ that lists the variable d-*loop* sites for several species and subspecies (see Nijman^25^). The last complication is the confusing nomenclatural changes. The authors proposed to replace the type species *P. femoralis* (Martin 1838) with a junior synonym (*P. neglectus neglectus* Schlegel 1876)^30^ without considering the detailed information in Low and Lim^31^ that explains why Martin is the author of the name *femoralis* and Singapore the type locality of the species. Abdul-Latiff et al.^28^’s study furthermore violated its own proposed nomenclatural changes by retaining *P. f. percura* and *P. f. robinsoni* (see Nijman^25^).

Here, we solve these problems by providing the first mitogenomes of *P. f. percura* and thus addressing the taxonomic position of all three subspecies of banded langurs. We also obtain the first mitogenome of the Riau pale-thighed langur *P. siamensis* cf. *cana* from Sumatra which helps with resolving species limits within this species. Lastly, we provide an updated dated phylogenetic tree for Asian colobines based on mitochondrial genomes and survey the mammal literature to illustrate that faecal DNA is currently still an underutilized source of genetic information.

## Results

### Survey of Zoological Record

In order to investigate to what extent faecal samples have been used in addressing taxonomic problems, we surveyed the literature as captured in Zoological Record. We retrieved 1,852 articles that mentioned faecal samples, but only a subset of 43 articles were also classified under Systematics/Taxonomy. Inspection of these records revealed only two studies that used faecal DNA for resolving species limits^32,33^.

### Sequence Data

Illumina sequencing of faecal metagenomes yielded 60.3-69.7 million sequences for each sample from Sumatra (ESBL1-8, Pres2; Table 2). The data were combined with the Hi-Seq data for six samples from Singapore (BLM1-6)^9^. All data were quality trimmed using Trimmomatic^34^ and complete mitochondrial genomes were obtained. One sample of *Presbytis femoralis percura* ESBL_7 had a low average coverage of <5X for the mitochondrial genome and was not analysed further.

**Table 2.** Summary of sequencing data and mitochondrial genomes obtained from eleven *Presbytis* langurs from Singapore and eastern Sumatra.

| Sample ID | Location | Organism | Raw/trimmed (millions) | Average mitochondrial coverage (X) |
| --- | --- | --- | --- | --- |
| ESBL 1b | Kampar | *Presbytis femoralis percura* | 60.33/58.81 | 25.205 |
| ESBL 5 | Bengkalis | *Presbytis femoralis percura* | 63.38/61.74 | 5.155 |
| ESBL 6a | Bengkalis | *Presbytis femoralis percura* | 67.04/66.61 | 8.721 |
| ESBL 8b | Bengkalis | *Presbytis femoralis percura* | 69.69/67.98 | 13.275 |
| Pres2 | Kampar | *Presbytis siamensis* cf. *cana* | 67.37/65.69 | 9.358 |
| BLM1 | Central Catchment Nature Reserve | *Presbytis femoralis femoralis* | 107.68/92.35 | 21.794 |
| BLM2 | Central Catchment Nature Reserve | *Presbytis femoralis femoralis* | 72.66/60.18 | 19.716 |
| BLM3 | Central Catchment Nature Reserve | *Presbytis femoralis femoralis* | 85.96/74.05 | 15.029 |
| BLM4 | Central Catchment Nature Reserve | *Presbytis femoralis femoralis* | 66.99/56.88 | 37.606 |
| BLM5 | Central Catchment Nature Reserve | *Presbytis femoralis femoralis* | 68.19/56.15 | 104.315 |
| BLM6 | Central Catchment Nature Reserve | *Presbytis femoralis femoralis* | 76.44/65.11 | 12.708 |

### Species Delimitation

Pairwise comparison of cyt-b, hypervariable region HV1 of the d-*loop* and mitochondrial genomes (CDS+rDNA+d-*loop*) revealed minimum genetic divergence of 7.1%, 6.1% and 5.3% between *P. f. femoralis* and *P. f. percura.* On the other hand, the minimum pairwise distance between either these taxa with *P. f. robinsoni* is 6.0% for HV1, 10.3% for cyt-b and 7.6% across the mitochondrial genome. For the two subspecies of *P. siamensis*, we were only able to compare HV1 sequences. The HV1 sequence of *P. s.* cf. *cana* is 11.1% diverged from *P. s. siamensis* and 5.1% from *P. melalophos* (KY117602), while cyt-b and complete mitochondrial genomes show divergence of 2.8% and 2.5% between sequences from *P. s.* cf. *cana* and *P. mitrata/P. melalophos.* Overall, these results suggest that *P. s.* cf. *cana* represents a genetically distinct *Presbytis* lineage.

The high genetic distances are consistent with results of species delimitation using Poisson Tree Processes (PTP)^35^, Automated Barcode Gap Discovery (ABGD)^36^ and Objective Clustering^37^. PTP consistently split *P. f. femoralis* and *P. f. percura* into different molecular Operational Taxonomic Units (mOTUs) across the three datasets examined (Asian colobine mitogenome and *Presbytis* mitogenome+cyt-b+HV1, and *Presbytis* HV1-only datasets) (Supplementary Figs. S1-3). ABGD and Objective Clustering (thresholds 2-4%) similarly split these two subspecies across different datasets (Asian colobine mitogenome, *Presbytis* HV1 and cyt-b) and a range of parameters (Table 3). For ABGD, these subspecies would only lump if unusually high priors for intraspecific divergences were used (priors>=0.0215). These parameters are not likely to be appropriate because they also led to collapse of many recognized *Presbytis* species into a single mOTU. All three species delimitation methods also placed *P. f. robinsoni* as a distinct species from *P. f. femoralis* and *P. f. percura.* Lastly, *P. s.* cf. *cana* was placed as a distinct species using PTP and ABGD unless inappropriately high priors for intraspecific divergence are used in ABGD. Objective Clustering based on HV1 also identified *P. s.* cf. *cana* as a distinct species. However, the mitogenome and cyt-b datasets lumped *P. s.* cf. *cana* with *P. melalophos* and *P. mitrata* at 3% and 4% thresholds.

**Table 3.**
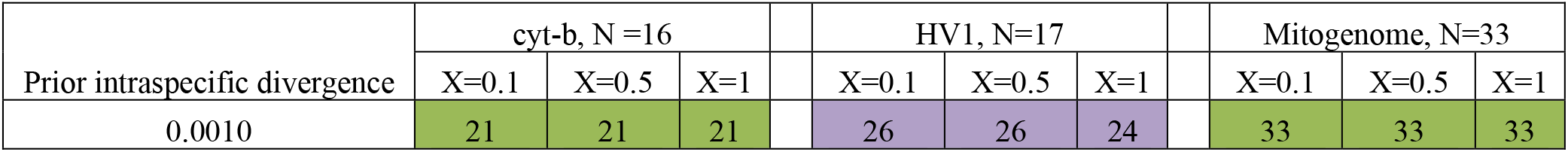
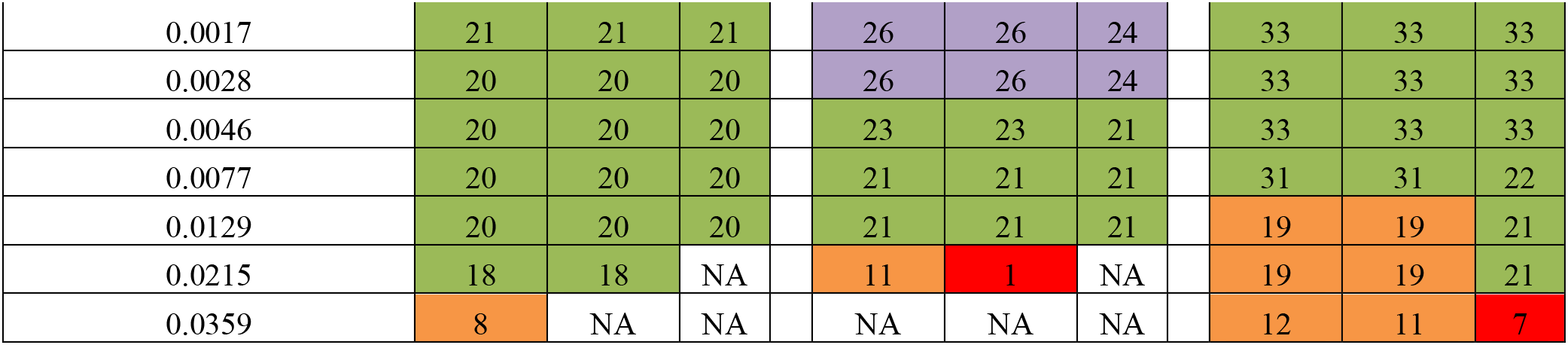
Summary of results of ABGD-based species delimitation. Values represent number of molecular Operational Taxonomic Units (mOTUs). N represents number of species (include those resurrected in this study), X the slope parameter. Colours: green: all resurrected species are valid. Purple: Supports splitting of the three subspecies of *Presbytis femoralis* and splitting of *P. siamensis* cf. *cana*. However, some sequences of *P. f. femoralis* from Peninsular Malaysia are split into multiple mOTUs. Orange: Resurrection of all three subspecies of *P. femoralis* is valid, but *P. s.* cf. *cana* is lumped with other *Presbytis* species. Red: Not valid.

We observed further species-level splitting when species delimitation was based on only HV1 data for the subspecies of *P. femoralis*. This dataset included sequences for multiple individuals of *P. f. femoralis, P. f. robinsoni* and *P. s. siamensis* from the Malay Peninsula (reconstructed in Nijman^25^, based on Abdul Latiff et al.^28^). At low prior intraspecific divergences, ABGD split some haplotypes of *P. f. femoralis* from the Malay Peninsula as separate mOTUs from other haplotypes of the same subspecies from the same region. These however consistently grouped together as single mOTU at higher thresholds. Similarly, PTP based analyses of only HV1 data split haplotypes of *P. f. robinsoni* into multiple mOTUs (Fig. S2) while ABGD consistently placed them as a single species.

### Mitochondrial Phylogeny of Asian Colobines and Genus Presbytis

The phylogenetic reconstruction based on mitochondrial genomes of Asian colobine dataset revealed that *P. femoralis* is polyphyletic (Fig. 3). The reconstructions based on Maximum Likelihood (ML) and Bayesian Inference (BI) are congruent and reveal that *P. f. femoralis* and *P. f. percura* are sister taxa. Divergence time estimates dated the split of *P. f. femoralis* and *P. f. percura* at 2.6 Mya (CI: partitioning by codon: 1.96-3.35 Mya, partitioning by gene 1.90-3.37 Mya (Supplementary Fig. S4)). This clade is sister to clade comprising of *P. mitrata, P. comata, P. siamensis* cf. *cana*, and *P. melalophos*. *Presbytis f. robinsoni* diverged from these species at 4.5 Mya (CI: partitioning by codon: 3.49-5.48 Mya, partitioning by gene 3.46-5.62). Overall, the mitochondrial phylogeny reveals high support for clade comprising of *Presbytis* and *Trachypithecus* as well as all the relationships within the (((*Simias+Nasalis*)*+Pygathrix*)*+Rhinopithecus*) clade. Only the placement of *Semnopithecus* remains uncertain, as revealed by low support for its relationship to the clade comprising of *Nasalis*, *Simias*, *Pygathrix* and *Rhinopithecus* on the ML tree. This result is different from Wang et al.^38^, who found high support for a sister group relationship of *Semnopithecus* to all the other genera of Asian colobines. However, a combined analysis of nuclear and mitochondrial data placed *Semnopithecus* differently thus suggesting our mt-genome phylogeny correctly reflects that the placement of *Semnopithecus* remains uncertain.

**Figure 3.**
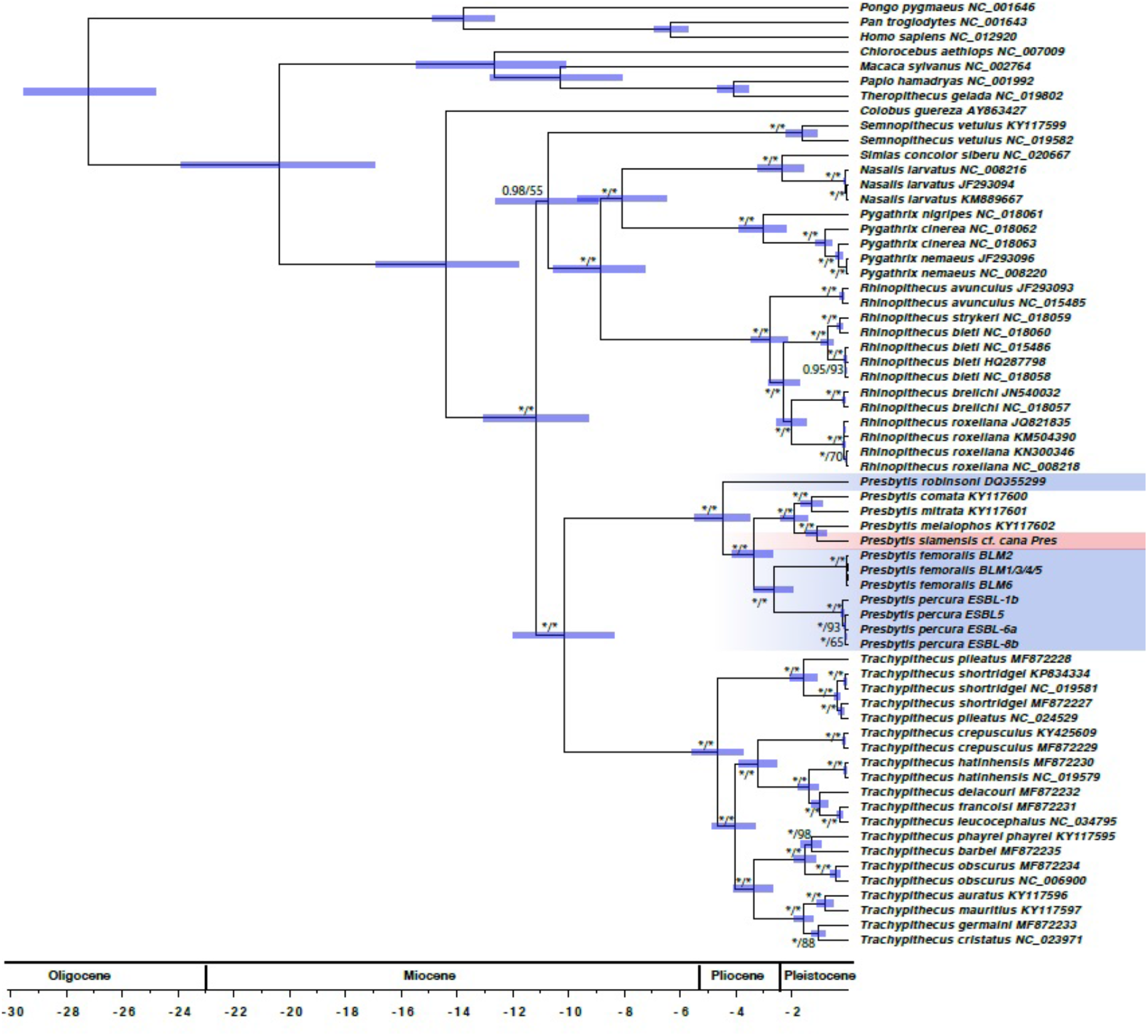
Dated phylogeny of Asian colobine primates based on mitochondrial genomes. The values at nodes represent posterior probability (codon partitioning)/ML bootstrap support for relationships between Asian colobines. Values are omitted if both BI and ML support values are <0.7/70, while * represents support of 1/100. The bars represent the 95% confidence intervals for divergence times estimates.

With regard to the phylogenetic relationships within *Presbytis*, our results on *Presbytis* mitogenome+cyt-b+HV1 dataset (Fig. 4) are largely consistent with the reconstruction by Meyer et al.^11^. The only differences are as follows: low support for a clade comprising of *P. comata, mitrata, melalophos, bicolor, sumatrana, rubicunda* and *P. siamensis* cf. *cana* but resolution for *P. rubicunda, melalophos, mitrata* and *bicolor* which formed a trichotomy in Meyer et al.^11^. Here, we found *P. rubicunda* to be sister to *P. bicolor. Presbytis f. femoralis* and *P. f. percura* remain sister taxa. Both taxa combined are more closely related to *P. mitrata, P. comata* and several other taxa of *Presbytis* than *P. f. robinsoni.* The split between *P. f. femoralis* and *P. f. percura* is again deeper than for most recognized taxa of *Presbytis.* Divergence estimates based on cyt-b for this taxon set revealed deeper divergence times as compared to mitochondrial genomes, but with overlapping confidence intervals. *Presbytis f. femoralis* and *P. f. percura* split 2.93 Mya (2.09-3.78 Mya) (Supplementary Fig. S5), while *P. f. robinsoni* diverged from the clade comprising of *P. femoralis, potenziani, mitrata, melalophos, bicolor, sumatrana* and *P. siamensis* cf. *cana* at 5.47 Mya (CI: 4.28-6.66 Mya).

**Figure 4.**
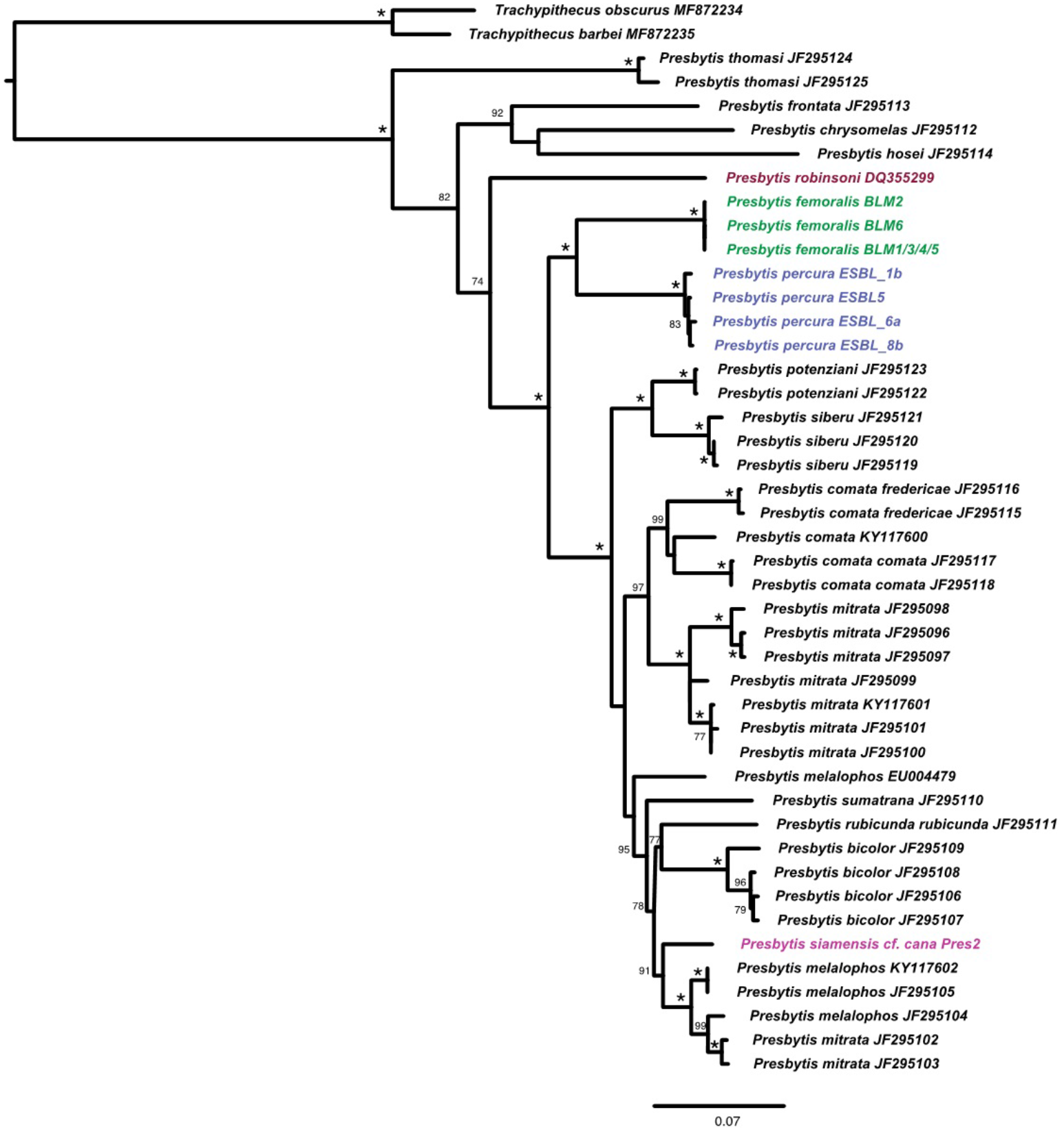
ML reconstruction of relationships between *Presbytis* species based on mitogenome+cyt-b+HV1 dataset. Node values represent bootstrap support, values <70 are excluded, while node support of 100 is represented by *.

## Discussion

The species limits of many Southeast Asian mammal taxa remain unclear which interferes with a timely conservation assessment although many populations, subspecies, and species face extinction. We here demonstrate how such taxonomic uncertainty can be addressed rapidly through shotgun sequencing of faecal DNA. We document the power of the approach by studying langur species in *Presbytis* Eschscholtz, 1821 which continue to undergo many changes that significantly affect the conservation status of many taxa. At one point all Asian langurs and leaf monkeys in *Presbytis*, *Semnopithecus*, and *Trachypithecus* were included in *Presbytis* and only five widespread species were recognized (*P. aygula*, *P. melalophos*, *P. frontata*, *P. potenziani*, *P. rubicunda*)^39,40,41^. This has dramatically changed over the last 20 years and currently three genera and 45 species are recognized (17 spp. in *Presbytis*; eight spp. in *Semnopithecus*; 20 spp. in *Trachypithecus*)^5,10^. Many of these changes in species boundaries were based on newly-obtained genetic data which allowed for the application of explicit species delimitation methods. These new data and analyses revealed that many taxa that were initially described as species and later downgraded to subspecies diverged well before the Pleistocene and should be recognized as species; i.e., the morphological characters that were used for the initial species descriptions were appropriate for the delimitation of species and the subsequent lumping was not justified.

### Resurrection of Presbytis femoralis, P. percura and P. robinsoni

Based on multiple species delimitation methods, high genetic divergence, placement in the mitochondrial phylogenies, as well as distinct morphological differences, we here resurrect the three species of *P. femoralis* from their current subspecific status (Table 4). The Raffles’ banded langur *P. femoralis* is only known from southern Peninsular Malaysia (states of Johor and Pahang) and Singapore. The East Sumatran banded langur *P. percura* only occurs in Riau Province of east-central Sumatra. Lastly, Robinson’s banded langur *P. robinsoni* has the widest distribution and ranges from northern Peninsular Malaysia (states of Kedah and Perak) through southern Thailand (provinces of Surat Thani, Phetchaburi, and Prachuap Khiri Khan) to southern Myanmar (Tanintharyi Region). These changes to species status mean that *Presbytis* now comprises 19 species (Fig. 5).

**Table 4.** Taxonomic classifications of the type specimens of *femoralis*, *percura*, and *robinsoni* followed by later authors since their first descriptions (non-exhaustive).

| Reference | *femoralis* | *percura* | *robinsoni* |
| --- | --- | --- | --- |
| Martin 1838 (16) | *Semnopithecus femoralis* | - | - |
| Lyon 1908 (18) | - | *Presbytis percura* | - |
| Thomas 1910 (19) | - | - | *Presbytis robinsoni* |
| Elliot 1913 (73) | *Pygathrix femoralis* | *Pygathrix percura* | *Pygathrix robinsoni* |
| Miller 1913 (74) | *Presbytis femoralis* | - | *Presbytis keatii* |
| Miller 1934 (72) | *Presbytis femoralis* | *Presbytis percura* | *Presbytis keatii* |
| Pocock 1934 (75) | *Presbytis femoralis femoralis* | *Presbytis melalophos percura* | *Presbytis femoralis keatii* |
| Raven 1935 (76) | *Presbytis femoralis* | *Presbytis percura* | *Presbytis robinsoni* |
| Chasen 1940 (77) | *Pithecus femoralis femoralis* | *Pithecus femoralis percura* | *Pithecus femoralis robinsoni* |
| Hooijer 1962 (78) | *Presbytis melalophos femoralis* | - | - |
| Medway 1970 (79) | *Presbytis melalophos femoralis* | - | *Presbytis melalophos robinsoni* |
| Thorington and Groves 1970 (80) | *Presbytis melalophos femoralis* | *Presbytis melalophos percura* | *Presbytis melalophos robinsoni* |
| Wilson and Wilson 1977 (81) | - | *Presbytis femoralis percura* | - |
| Medway 1983 (82) | *Presbytis melalophos femoralis* | - | *Presbytis melalophos robinsoni* |
| Brandon-Jones 1984 (83) | *Presbytis femoralis femoralis* | *Presbytis femoralis percura* | *Presbytis femoralis robinsoni* |
| Napier 1985 (84) | *Presbytis melalophos femoralis* | *Presbytis melalophos percura* | *Presbytis melalophos robinsoni* |
| Weitzel et al. 1988 (21) | *Presbytis femoralis femoralis* | - | *Presbytis femoralis robinsoni* |
| Aimi and Bakar 1992 (85) | - | *Presbytis femoralis percura* | - |
| Oates et al. 1994 (12) | *Presbytis melalophos femoralis* | *Presbytis melalophos percura* | *Presbytis melalophos robinsoni* |
| Groves et al. 2001 (15) | *Presbytis femoralis femoralis* | *Presbytis femoralis percura* | *Presbytis femoralis robinsoni* |
| Brandon-Jones et al. 2004 (86) | *Presbytis femoralis femoralis* | *Presbytis femoralis percura* | *Presbytis femoralis robinsoni* |
| Md.-Zain 2001 (87) | *Presbytis melalophos femoralis* | - | *Presbytis melalophos robinsoni* |
| Meyer et al. 2011 (11) | *Presbytis femoralis femoralis* | - | *Presbytis femoralis robinsoni* |
| Vun et al. 2011 (27) | *Presbytis melalophos femoralis* | - | *Presbytis melalophos robinsoni* |
| Roos et al. 2014 (10) | *Presbytis femoralis femoralis* | *Presbytis femoralis percura* | *Presbytis femoralis robinsoni* |
| Abdul Latiff et al. 2019 (28) | *Presbytis neglectus neglectus* | *Presbytis femoralis percura* | *Presbytis femoralis robinsoni* |
| This study | *Presbytis femoralis* | *Presbytis percura* | *Presbytis robinsoni* |

Grey boxes indicate the recognition of species rank for *Presbytis femoralis*, *P. percura*, and *P. robinsoni*.

**Figure 5.**
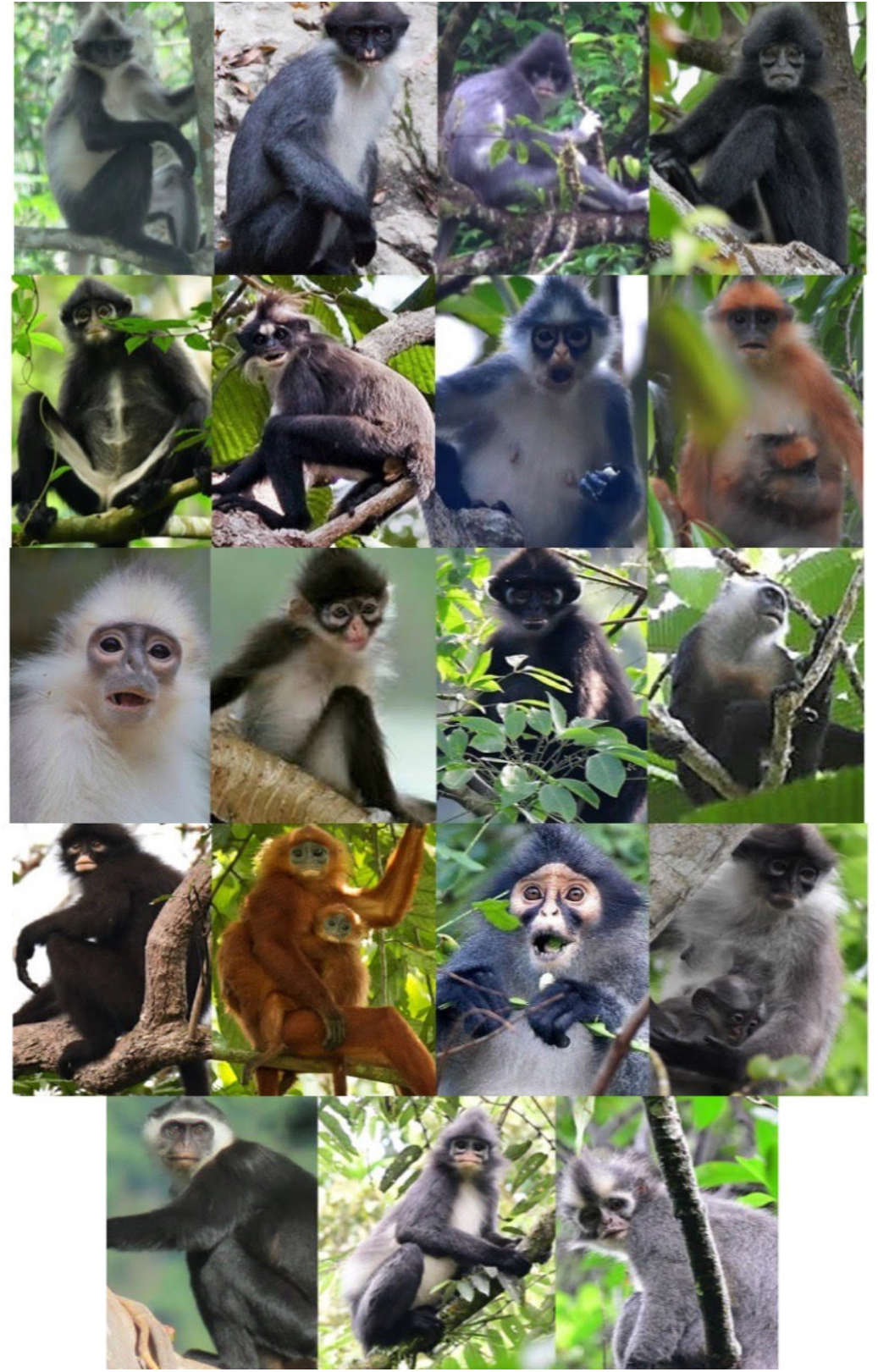
Nineteen species of *Presbytis* langurs recognised in this study. Photo credits: Wilson Novarino (1. *P. bicolor*); Brent Loken (2. *P. canicrus*); Chien Lee (3. *P. chrysomelas*); Andie Ang (4. *P. comata*); Andie Ang (5. *P. femoralis*); Milan Janda (6. *P. frontata*); Mark Spence (7. *P. hosei*); Muhammad Azhari Akbar (8. *P. melalophos*); Jonathan Beilby (9. *P. mitrata*); Martjan Lammertink (10. *P. natunae*); Andie Ang (11. *P. percura*); SwaraOwa (12. *P. potenziani*); Andie Ang (13. *P. robinsoni*); David Tan (14. *P. rubicunda*); Coke and Som Smith (15. *P. sabana*); Lee Zan Hui (16. *P. siamensis*); Gusmardi Indra (17. *P. siberu*); Thio Hui Bing (18. *P. sumatrana*); Thio Hui Bing (19. *P. thomasi*). Collage credit: Sabrina Jabbar

A >5% genetic difference and divergence estimates of 2.6-2.9 Mya between *P. femoralis* and *P. percura* suggest that these two species radiated prior to the Pleistocene, while several other species of *Presbytis* diverged more recently. These results are particularly intriguing because the changing sea levels during Pleistocene would have increased connectivity between the land masses of Sumatra and Malay Peninsula. However, the Malacca Straits River flowing northwards with tributaries in what is now Sumatra and the Malay Peninsula^42^ may have been a substantial barrier between *P. femoralis* and *P. percura*, as it would have been significantly wider than the rivers that currently form the geographic barriers between some of the *Presbytis* species in Sumatra. Furthermore, it has been argued that the land bridge had coarse sandy and/or poorly drained soils which may have limited plant growth in central Sundaland. Unsuitable vegetation may have acted as a dispersal barrier for rainforest plants and animals^43^. These barriers could have kept the langur populations separate and it thus remains unclear whether *P. femoralis* and *P. percura* would have formed a hybrid population if they had encountered each other. Note that it is known that recognized primate species in *Trachypithecus* that radiated ~0.95-1.25 Mya can interbreed^44,45^, but the genetic divergence between *P. femoralis* and *P. percura* is considerably higher.

### Conservation Status of P. femoralis, P. percura, and P. robinsoni

In the most recent IUCN Red List assessment conducted in 2015, *Presbytis femoralis* (comprising *femoralis*, *percura* and *robinsoni*) was listed as Vulnerable A2cd A3cd A4cd (population size reduction of at least 30% over three generations based on a decline in area of occupancy, extent of occurrence and habitat quality, and actual or potential levels of exploitation). As part of this assessment the status of the three subspecies were also evaluated against the IUCN Red List criteria. *Presbytis f. femoralis* was considered Endangered A2cd A3cd A4cd, *P. f. percura* Data Deficient, and *P. f. robinsoni* Near Threatened (unpublished data from a Red List re-assessment in 2015). With their resurrection to species rank, the conservation status of each of the taxa requires re-assessment. *Presbytis femoralis* has a small global population size which continues to decline mainly due to habitat loss. There are 60 individuals (48 mature individuals) in the Singapore population of *P. femoralis*^46^. There are no precise population estimates available for the conspecifics in the Malaysian states of Johor and Pahang, but it is believed that only a few hundred individuals remain (see Abdul-Latiff et al.^28^); i.e., the overall population of *P. femoralis* could well be <250 mature individuals.

Furthermore, the extensive habitat loss especially to industrial oil palm plantations in southern Peninsular Malaysia is unlikely to cease in the near future (see Shevade et al.^47^; Shevade and Loboda^48^). Hence, based on a small population size and decline, we propose to list *P. femoralis* as Critically Endangered C2a(i) (<250 mature individuals, continuing population decline, and ≤50 mature individuals in each subpopulation).

*Presbytis percura* is only found in a number of isolated forests and faces extinction in the wild based on large-scale forest loss in Riau Province^49^. Riau experienced the highest rate of deforestation in Sumatra such that 63% of natural forest have been lost between 1985 and 2008^50^. Additionally, forest fires linked to the ENSO events, and open burning of forest land for agricultural purposes destroy millions of hectares of land in Indonesia on an annual basis, and Riau is often one of the worst impacted areas, owing in part to its high concentration of peatland^51^. We thus infer that the area of occupancy, extent of occurrence and quality of habitat of *P. percura* have declined such that their population size has reduced by ≥80% over the last three generations since 1989 (30 years approximately; see Nijman and Manullang^52^ for the closely-related *P. melalophos*), thus fulfilling the IUCN criteria for Critically Endangered A2cd A3cd A4cd.

*Presbytis robinsoni* ranges from northern Peninsular Malaysia through southern Thailand to southern Myanmar. There are no population estimates (either recent or in the past), but some of the species’ habitat continues to be converted for agriculture (primarily oil palm) and it is also targeted for illegal pet trade. Overall, it cannot be evaluated on small population size and/or restricted population criteria, but *P. robinsoni* is certainly a taxon of conservation concern and is here considered as Near Threatened.

### An Urgent Need for Molecular Data for Additional Presbytis Populations

Our results highlight the need for sampling multiple populations of *Presbytis* species and subspecies because even our limited fieldwork already provided strong evidence for the widespread presence of cryptic diversity or inappropriate synonymization within *Presbytis.* Additional data are also needed in order to be able to precisely assign samples. We collected one faecal sample that was suspected to come from an individual of *P. siamensis* cf. *cana.* However, its placement in the phylogeny reveals that it belongs to a genetically distinct lineage that is more closely related to *P. melalophos+mitrata* than *P. s. siamensis* (Supplementary Fig. S2). If the sample was indeed from *P. s. cana*, then the taxonomy of the pale-thighed langur *P. siamensis,* which currently comprises of four subspecies, needs to be revisited unless the unexpected signal is due to introgression via the hybridization of two species. Regardless of the explanation, the taxon represented by the sample deserves species status, but the correct scientific name and range limits remain unclear because *P. s. paenulata* and *P. s. rhionis* still lack molecular data. Even if the faecal sample originated from individual of *P. melalophos/mitrata*, its genetic distinctness suggests that these species require more attention from taxonomists. In addition, it would mean that the geographic ranges of the species need to be revised because the species are unknown from the place of collection. Overall, either explanation is reasons for concern. Geographically, only *P. s. siamensis* (Fig. 6; left photo) has a wide distribution on the Malay Peninsula while the remaining three subspecies have narrow distributions. *Presbytis s. cana* (Fig. 6; right photo) occurs in eastern Sumatra and on Kundur Island, *P. s. paenulata* is found mainly in a small-wedge of coastal forest in east-central Sumatra, and *P. s. rhionis* has only been found on the islands of Bintan and Batam (but may also be found on Galang Island) in the Riau Archipelago^10^. Given the extensive habitat loss to oil palm plantations in Sumatra and large-scale economic development in Bintan and Batam, these taxa are likely highly threatened. Study is urgently needed and we submit that faecal samples would be the best way to rapidly address the species limits and distribution of these undersampled *Presbytis* species. We lack genetic data, even COI barcodes, for many subspecies of *Presbytis* and their distributions are poorly understood. Clearly, Southeast Asian langurs have received insufficient attention and broad surveys are needed that estimate population sizes while collecting faecal samples for a re-assessment of species boundaries

**Figure 6.**
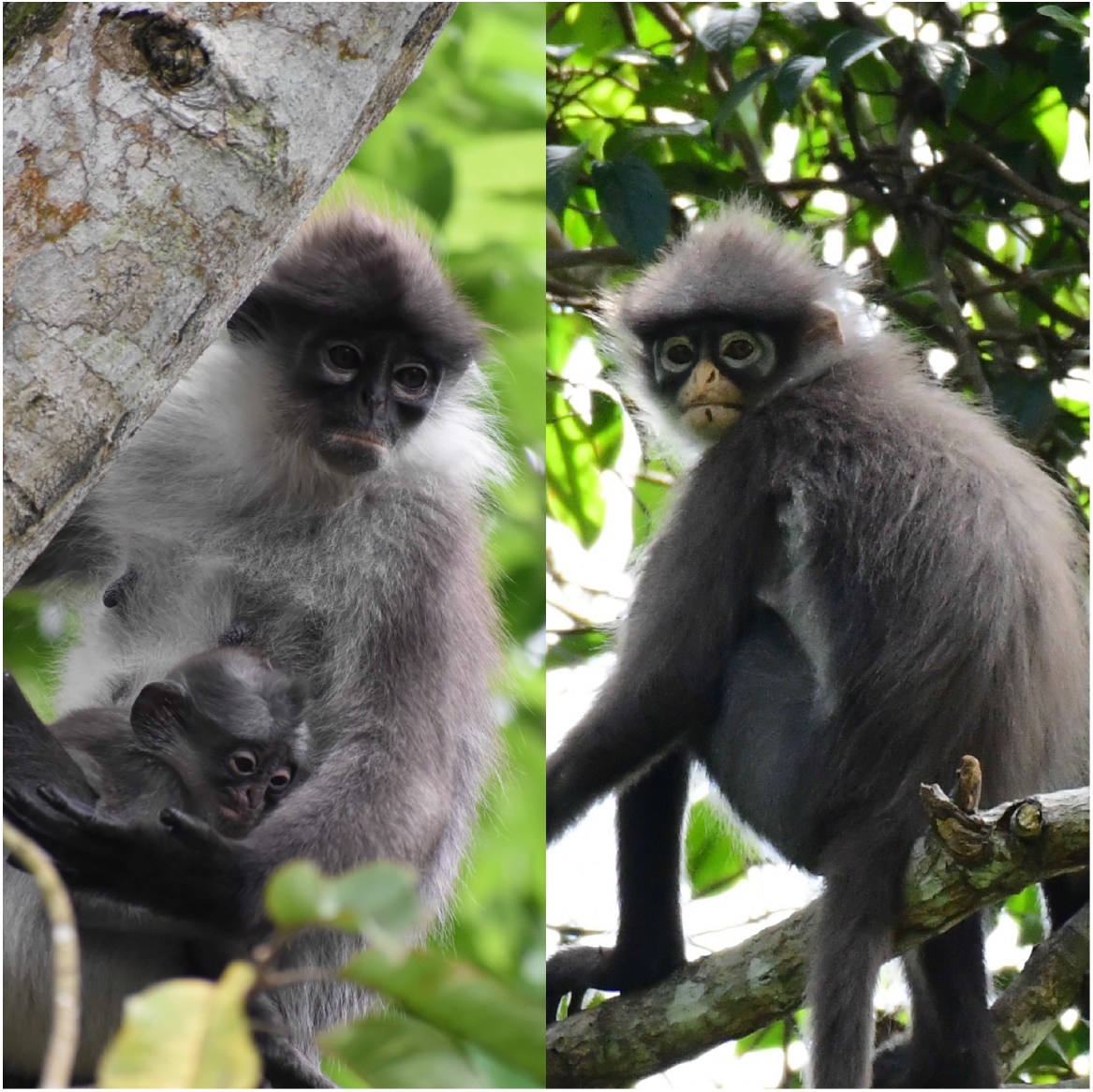
*Presbytis siamensis siamensis* (left; photo: Lee Zan Hui) and *P. s. cana* (right: Andie Ang).

### Mitochondrial Phylogeny of Presbytis

In the process of delimiting species, we re-examined the phylogenetic relationships within *Presbytis* by combining the data for mitochondrial genomes with the data generated by Meyer et al.^11^ for cyt-b and d-*loop*-HV1. This led to the resurrection of *P. femoralis* and *P. percura* which are here revealed to be sister species and more closely related to several other *Presbytis* species from Sumatra (and *P. rubicunda* in Borneo) than *P. robinsoni* from Malay Peninsula. This placement of *P. femoralis* is in conflict with relationships proposed by Abdul-Latiff et al.^28^, who obtained a clade comprising of *P. femoralis* + *P. robinsoni* + *P. siamensis* which was sister species of the *Presbytis* species from Sumatra + *P. rubicunda*.

One limitation of our study is the lack of nuclear data for species delimitation and reconstructing relationships (e.g. Wang et al.^38^). This is because mitochondrial data represents matrilineage only. However, obtaining nuclear data from faecal samples remains challenging partially due to the low concentration of primate DNA in faecal samples. We assessed the primate nuclear DNA content in these metagenomes and found that it was only 0.09-3.13% of total DNA in the faecal samples (Table 5). However, we do not think that the lack of nuclear data seriously challenges our conclusions. Firstly, we reveal deep mitochondrial splits of >2 million years between *P. femoralis* and *P. percura*. We think that it would be more perilous to argue for the presence of one species based on identical mitochondrial sequence because it could be caused by introgression. Secondly, these splits are consistent with morphological data that allow for assigning specimens unambiguously to one of the lineages that are here re-corrected as species. Lastly, whatever limited information is available for Asian colobines does not point to widespread conflict between nuclear and mitochondrial signals. Wang et al.^38^ presents phylogenetic reconstructions based on mitochondrial and nuclear markers for Asian colobines. Four of the six congeneric nodes are congruent between nuclear and mitochondrial data although they came from different individuals (i.e., study used data from multiple sources). Nonetheless, it is essential to develop new approaches for obtaining nuclear data from faecal samples (see Chiou and Bergey^53^).

**Table 5.** Estimation of host DNA content in faecal metagenomes

| Sample ID | % host | % host after removal of human reads |
| --- | --- | --- |
| ESBL 1b | 0.51 | 0.49 |
| ESBL 5 | 0.62 | 0.59 |
| ESBL 6a | 0.23 | 0.21 |
| ESBL 7 | 2.00 | 1.95 |
| ESBL 8b | 3.15 | 3.13 |
| Pres 2 | 0.11 | 0.09 |

### Non-invasive Samples for Species Discovery

Seventeen (61%) of the 28 taxa of *Presbytis* are threatened (Vulnerable, Endangered, or Critically Endangered) while five (18%) taxa are Data Deficient^5,10^. Many taxa continue to be affected by habitat loss. This means that there is an urgency to resolve species limits in order to better assess their conservation status and needs. Molecular data play a critical role in resolving species limits, and here faecal samples are particularly valuable because they allow for accelerating data collection. Yet, our literature survey suggests that faecal DNA remains underutilized for taxonomic research with only two other published studies explicitly using faecal DNA as evidence for justifying decision on species status: nine members from a brown lemur complex were given species rank^33^ and two subspecies of otters were elevated to species^32^. Faecal samples yield DNA that are valuable not only for taxonomic purposes but also population genetics, diet analyses, microbiome and parasite research, and should be routinely collected during field surveys. In groups such as *Presbytis,* faecal samples would be particularly useful as these animals are shy and many populations are highly threatened.

## Conclusions

We here demonstrate the value of non-invasive faecal samples for addressing taxonomic questions that are of significant conservation importance. Based on mitochondrial DNA (mitogenomes, cyt-b and d-*loop*), we resurrected three species within the *Presbytis femoralis* group. The new species limits also led to a change in the conservation status of *P. femoralis* and *P. percura* which now have to be considered Critically Endangered. We further urge researchers to include the collection of non-invasive faecal samples into their field protocols.

## Materials and Methods

### Literature Survey

In order to assess how frequently faecal samples are used for addressing taxonomic questions, we conducted a survey of mammalian literature for the last 41 years (between 1978-2018). We downloaded the records pertaining to Supertaxon Mammalia (ST=Mammalia) from *Zoological Record* (articles only). We then identified a subset containing valid mammalian species names by using the checklist by Burgin et al.^54^. This was done by searching the binomial name of 6,399 species in the list. We also searched for the genus and species names separately to include records that utilize genus abbreviations. We then retrieved the studies that involved DNA/molecular work on faecal samples by searching for the terms (feces/faeces/fecal/faecal/scat/scats) and (DNA/barcod/sequenc/molecul/genom/genetic/microsatellite). We lastly retrieved the records that were classified under Systematics/Taxonomy in the “Broad terms” field and examined the records manually.

### Sample Collection, DNA Extraction, and Sequencing

Five faecal samples of *Presbytis femoralis percura* were collected during an eight-day (25 April - 2 May 2018) survey in Riau Province^49^. We also collected one faecal sample believed to come from *P. siamensis cana* (herein referred as *P. siamensis* cf. *cana*) as the monkeys were seen on the same tree below which the fresh faecal sample was found. The samples were preserved following a two-step ethanol-silica method^55^ and subsequently stored at a −20°C freezer at the Andalas University in Sumatra. Genomic DNA was extracted at Andalas University from 50 mg of faeces using QIAamp^®^ Fast DNA Stool Mini Kit (QIAGEN, Singapore). DNA was recovered in 30 μl of elution buffer (instead of 200 μl) in order to obtain a higher concentration of DNA. Each sample was also extracted 2-3 times and later pooled to recover more genomic DNA. Genomic DNA of these six samples were sent from Andalas University for Illumina HiSeq 4000 (Illumina Inc., San Diego, CA) sequencing (150 PE) by a commercial provider (NovogeneAIT). A library was constructed for each faecal sample (fragment size 350 bp) using NEBNext Ultra II DNA Library Prep Kit.

### Bioinformatics for Obtaining Mitochondrial Genomes

Raw reads generated for *P. f. percura* and *P. s. cana* in this study as well as those for *P. f. femoralis* from Srivathsan et al.^9^ were trimmed using Trimmomatic v.0.33^34^ under the following parameters: LEADING:3 TRAILING:3 SLIDINGWINDOW:4:15 MINLEN:50, and the ILLUMINACLIP parameter was set at 2:30:10^34^. New mitochondrial reference genomes were assembled for *P. f. percura and P. s. cana* using MITObim v.1.9^56^ using the available mitogenome of *P. f. femoralis* (KU899140) from one sample each (ESBL1b and Pres2). Any redundancy due to circular nature of mitochondrial genome was removed in the resulting assembly. We then obtained mitochondrial genomes per faecal sample by mapping the quality-trimmed reads for the sample to the reference genome (ESBL samples to mitochondrial genome from ESBL1b, Pres2 to Pres2 and BLM1-6 to KU899140). Reads were mapped using Bowtie2^57^ under paired end and -end-to-end mode. The resulting SAM files were converted to BAM files using SAMtools^58^. Variants were detected using Lofreq^59^ using parameters similar to Isokallio and Stewart^60^ with a few more stringent criteria: we included mapping quality in LoFreq’s model and also retained base-alignment quality. We furthermore applied a minimum allele frequency of 0.2 while filtering the vcf to accept heteroplasmy only if it was at a frequency of >0.2. Alternate mitochondrial genomes were reconstructed from the resulting vcf file using gatk^61^ FastaAlternateReferenceMaker and heteroplasmic sites were modified as ambiguous nucleotides using custom script. All mitochondrial genomes were annotated using MITOS^62^.

### Phylogenetic Reconstructions and Species Delimitations

The newly reconstructed mitochondrial genomes were combined with publicly available data from GenBank. For the latter, we downloaded all the mitochondrial sequences for colobine primates and then retained only the data for Asian colobines. We then curated the GenBank records by consulting the source publication and assessing the locality information provided in order to update the taxonomic names given that many subspecies are now considered species. We excluded those sequences for which the source information was incomplete. This curated set of sequences was used for downstream distance based (Automated Barcode Gap Discovery, ABGD^36^ and Objective Clustering^37^ and tree-based species delimitation analyses (Poisson Tree Processes or PTP)^35^. We also included data for d-*loop* HV1 obtained by Abdul-Latiff et al.^28^ for *P. f. femoralis, P. f. robinsoni* and *P. siamensis siamensis* from Malaysia. Given that these sequences were not submitted to GenBank, we used Nijman^25^’s reconstruction of the sequences based on a table that lists all variable sites relative to a reference sequence.

For analyses with PTP, we used three datasets: (1) The Asian colobine mitogenome dataset based on genomes for Asian colobines (minimum length >10,000 bp). The sequences for the 13 mitochondrial CDS, two ribosomal genes, and complete d-*loop* sequences were extracted, aligned, and concatenated. Here, *Colobus guereza* and *Macaca sylvanus* were selected as the outgroups. (2) A second dataset included the *Presbytis* mitogenome+cyt-b+HV1 dataset. This dataset covers more samples because *Presbytis* has been well sampled for cyt-b and the hypervariable region I (HV1) of d-*loop* (see Meyer et al.^11^). For the analyses of this dataset, we used *Trachypithecus obscurus* and *T. barbei* as outgroups. (3) The last dataset comprised HV1 sequences only (*Presbytis* HV1-only dataset). This included HV1 sequences obtained by Abdul-Latiff et al.^28^ for *P. f. femoralis, P. f. robinsoni* and *P. s. siamensis*.

All coding sequences were aligned in MEGA X^63^ based on amino acid translations (using Clustal). The ribosomal genes and d-*loop* sequences were aligned using MAFFT LINSI^64^. We ensured that only distinct haplotypes were retained and only retained the longest sequence if identical sequences were found. The alignments were concatenated in SequenceMatrix 1.7.8^65^. Maximum Likelihood reconstructions were carried out using RAxML v8^66^. For the mitogenome and the *Presbytis* mitogenome+cyt-b+HV1 datasets, we determined best partitioning scheme by providing 42 different partitions to PartitionFinder^67^ corresponding to codon position for the 13 coding regions and 3 separate partitions for 12S, 16S and d-*loop*. RAxML was run using GTRGAMMA with the resulting partitioning scheme (no partitioning was done for the HV1 dataset) with 20 independent searches for the best tree. Multiparametric bootstrapping was conducted applying the automatic bootstopping criterion (autoMRE). The resulting trees were subjected to PTP-based species delimitation after excluding outgroups.

Distance-based species delimitation utilized ABGD based on uncorrected distances^68^. We assessed species delimitations under different parameters by varying the slope X=0.1, X=0.5 and X=1 (X=1.5 was not applicable to the dataset) and prior intraspecific divergences. Species delimitation was carried out based on the Asian colobine mitogenome dataset described above and the *Presbytis* cyt-b and HV1 alignments. The same datasets were also clustered using Objective Clustering as implemented in Species Identifier (Taxon DNA 1.6.2)^37^ at genetic distances of 2.0, 3.0 and 4.0%.

### Divergence Dating

Divergence dates between Asian colobine lineages were determined using BEAST v 2.6.0^69^. We used Asian colobine mitogenome dataset but excluded d-*loop* for this analyse due its differing mutational patterns as done for previous studies^11,70^. For divergence estimates, we included the following genomes to the Asian colobine mitogenome dataset: *Pongo pygmaeus* (NC_001646), *Pan troglodytes* (NC_001643), *Homo sapiens* (NC_012920), *Chlorocebus aethiops* (NC_007009), *Macaca sylvanus* (NC_002764), *Papio hamadryas* (NC_001992), *Theropithecus gelada* (NC_019802). We also did a second analysis for *Presbytis* cyt-b as done by Meyer et al.^11^ and used a similar strategy for the various steps. Here, representative sequences from different Asian colobine genera were included: *Trachypithecus obscurus* (NC_006900), *Nasalis larvatus* (NC_008216), *Rhinopithecus avunculus* (NC_015485), *Semnopithecus vetulus* (NC_019582) in addition to the above-mentioned sequences for fossil-based calibration. The fossil calibration dates used in this study also followed the dates used by Meyer et al.^11^. For mitochondrial genomes, we analysed the data using the following partitioning schemes: by codon (5 partitions: 1,2,3 codon for coding genes, 12S, 16S) and by gene. We also tested partitioning by both gene and codon, but found the Effective Sample Size (ESS) to be low for multiple parameters. For cyt-b dataset, we used the 1+2 and 3 codon partitioning scheme^11^.

Divergence estimates were based on a relaxed log normal clock and a Yule prior. Site models were unlinked across partitions, and a model-averaging approach was used as implemented in bModelTest^71^. Two independent runs were conducted with 25 million generations with sampling at every 1000 generations. Tracer v 1.7.1 was used to assess convergence, LogCombiner 2.6.1 was used to combine the results and 10% burn-in removal was applied. TreeAnnotator v 2.6.0 was used to summarize the trees.

### Estimation of Host DNA Content in Faecal Metagenomes

In order to estimate the amount of host nuclear DNA in the faecal metagenomes, we mapped the metagenomic reads to a colobine reference genome (*Rhinopithecus roxellana*: GCF_000769185.1) using bowtie-2 under --end-to-end and ---very-sensitive mode. We next excluded potentially contaminated reads that could correspond to humans. For this, the mapped reads were retrieved from the resulting bam files using samtools. These were mapped back to a combined reference dataset of *R. roxellana* and human genome (GCF_000001405.39, GRCh38) using Bowtie2. Resulting BAM file was filtered to exclude reads with any mismatches, and we then excluded all sequences that matched to human genome only.

### Ethics Statement

We followed the Code of Best Practices for Field Primatology (2014). All genetic material were obtained non-invasively through faecal samples; no animals were harmed in the process. We followed the rules and regulations of the Government of Indonesia and RISTEKDIKTI (research permit no. 3051/FRP/E5/Dit.KI/IX/2018).

## Acknowledgements

This research was funded by the Wildlife Reserves Singapore Conservation Fund. We would like to thank the Government of Indonesia and RISTEKDIKTI for the research permit (3051/FRP/E5/Dit.KI/IX/2018) and the Government of Singapore and National Parks Board for the research permit (NP/RP16-092). We sincerely appreciate the assistance from Anugrah Viona Agesi, Wila Karlina, and Dyta Rabbani Aidil of the Genetic and Biomolecular Laboratory at Andalas University.

## Author Contributions

A.A., A.S., R.M., V.N. wrote the main manuscript text; A.A., A.S., R., R.M., V.N. designed the study; A.A., R. conducted the field work; A.A., D.R. extracted DNA samples in D.R.’s laboratory; A.S., D.R. conducted bioinformatics analyses; and A.S. reconstructed the time-tree. All authors reviewed and edited the manuscript and gave final approval for submission.

## Additional Information

The authors declare no competing interests.

## References

1. IPBES. Summary for policymakers of the global assessment report on biodiversity and ecosystem services of the Intergovernmental Science-Policy Platform on Biodiversity and Ecosystem Services. in The IPBES Global Assessment on Biodiversity and Ecosystem Services (ed. Díaz, S. et al.) (IPBES secretariat, Bonn, Germany (2019).

2. Sodhi, N. S., Koh, L. P., Brook, B. W. & Ng, P. K. L. Southeast Asian biodiversity: an impending disaster. Trends Ecol. Evol. 19(12),654–660 (2004).

3. Trimble, M. J. & van Aarde, R. J. Geographical and taxonomic biases in research on biodiversity in human-modified landscapes. Ecosphere 3(12),1–16 (2012).

4. Titley, M. A., Snaddon, J. L. & Turner, E. C. Scientific research on animal biodiversity is systematically biased towards vertebrates and temperate regions. PLoS One 12(12),e0189577 (2017).

5. Mittermeier, R. A., Rylands, A. B. & Wilson, D. E. Handbook of the Mammals of the World, Vol. 3, Primates. Lynx Edicions, Barcelona, Spain (2013).

6. Mace, G. The role of taxonomy in species conservation. Phil. Trans. R. Soc. Lond. B Biol. Sci. 359(1444),711–719 (2004).

7. Thomson, S. A. et al. Taxonomy based on science is necessary for global conservation. PLoS Biol. 16(3),e2005075 (2018).

8. Srivathsan, A., Sha, J., Vogler, A. P. & Meier, R. Comparing the effectiveness of metagenomics and metabarcoding for diet analysis of a leaf‐feeding monkey (*Pygathrix nemaeus*). Mol. Ecol. Res. 15(2),250–261 (2015).

9. Srivathsan, A., Ang, A., Vogler, A. P. & Meier, R. Fecal metagenomics for the simultaneous assessment of diet, parasites, and population genetics of an understudied primate. Front. Zool. 13, 17 (2016).

10. Roos, C. et al. An updated taxonomy and conservation status review of Asian primates. Asian Primates J. 4(1),2–38 (2014).

11. Meyer, D. et al. Mitochondrial phylogeny of leaf monkeys with implications for taxonomy and conservation. Mol. Phylogenet. Evol. 59(2),311–319 (2011).

12. Oates, J. F., Davies, A. G. & Delson, E. The diversity of living colobines. In: Davies, A. G. & Oates, J. F. (eds). Colobine Monkey: Their Ecology, Behaviour and Evolution. Cambridge University Press (1994).

13. Nijman, V. Ecology of sympatric and allopatric *Presbytis* and *Trachypithecus* langurs in Sundaland. in The Colobines: Natural History, Behaviour and Ecological Diversity. (eds. Matsuda, I., Grueter, C. C. & Teichroeb, J. E). (Cambridge University Press, Cambridge, in press).

14. IUCN. The IUCN Red List of Threatened Species. https://www.iucnredlist.org (2008).

15. Groves, C. P. Primate Taxonomy. Smithsonian Institution Press, Washington (2001).

16. Martin, W. C. L. A monograph of the genus *Semnopithecus* (continued from page 326). Mag. Nat. Hist. 2, 434–441 (1838).

17. Raffles, S. T. Descriptive catalogue of a zoological collection made in the island of Sumatra and its vicinity. Trans. Linn. Soc. Lond. 13, 247 (1821).

18. Lyon, M. W. Jr. Mammals collected in eastern Sumatra by Dr. W. L. Abbott during 1903, 1906, and 1907: with descriptions on new species and subspecies. Proc. US Natl. Mus. 34, 619–679 (1908).

19. Thomas, O. A new monkey from the Malay Peninsula. Proc. Zool. Soc. Lond. (Abstr. p. 26), 634–635 (1910).

20. Robinson, H. C. & Kloss, C. B. On six new mammals from the Malay Peninsula and adjacent islands. J. Fed. Malay States Mus. 4(2),169–174 (1911)

21. Weitzel, V., Yang, C. M. & Groves, C. P. A catalogue of primates in the Singapore zoological reference collection. Raffles B. Zool. 36, 1–166 (1988).

22. Ang, A., Srivathsan, A., Md.-Zain, B. M., Ismail, M. R.B. & Meier, R. Low genetic variability in the recovering urban banded leaf monkey population of Singapore. Raffles B. Zool. 60, 589–594 (2012).

23. Ang, A. and Boonratana, R. *Presbytis femoralis* ssp. percura. In: The IUCN Red List of Threatened Species 2015 (in press).

24. Ang, A. Banded Leaf Monkeys in Singapore: Preliminary Data on Taxonomy, Feeding Ecology, Reproduction, and Population Size. MSc thesis, National University of Singapore (2010).

25. Nijman, V. *Presbytis neglectus* or *P. femoralis*, colobine molecular phylogenies, and GenBank submission of newly generated DNA sequences. Folia Primatol. (2019b).

26. Md.-Zain, B. M., Morales, J. C., Hassan, M. N., Jasmi, A. & Melnick, D. J. Is *Presbytis* a distinct monophyletic genus: inferences from mitochondrial DNA sequences. Asian Primates J. 1, 26–36 (2008).

27. Vun, V. F., Mahani, M. C., Lakim, M., Ampeng, A. & Md.-Zain, B. M. Phylogenetic relationships of leaf monkeys (*Presbytis*; Colobinae) based on cytochrome b and 12S rRNA genes. Genet Mol Res 10(1),368–381 (2011).

28. Abdul-Latiff, M. A. B., Baharuddin, H., Abdul-Patah, P. & Md.-Zain, B. M. Is Malaysia’s banded langur, *Presbytis femoralis femoralis*, actually *Presbytis neglectus neglectus*? Taxonomic revision with new insights on the radiation history of the *Presbytis* species group in Southeast Asia. Primates 60(1),63–79 (2019).

29. Sterner, K. N., Raaum, R. L., Zhang, Y.-P., Stewart, C.-B. & Disotell, T. R. Mitochondrial data support an odd-nosed colobine clade. Mol. Phylogenet. Evol. 40(1),1–7 (2006).

30. Schlegel, H. Monographie 40: simiae. Revue Methodique, Museum d’Histoire Naturelle des Pays-Bas 7, 1–356 (1876).

31. Low, M. E. Y. & Lim, K. K. P. The authorship and type locality of the banded leaf monkey, *Presbytis femoralis*. Nature Singapore 8, 69–71 (2015).

32. Koepfli, K.-P. et al. Establishing the foundation for an applied molecular taxonomy of otters in Southeast Asia. Conserv. Genet. 9, 1589 (2008).

33. Markolf, M. et al. True lemurs … true species - species delimitation using multiple data sources in the brown lemur complex. BMC Evol. Biol. 13, 233 (2013).

34. Bogler, A. M., Lohse, M. & Usadel, B. Trimmomatic: A flexible trimmer for Illumina Sequence Data. Bioinformatics 30, 2114–2120 (2014).

35. Zhang, J., Kapli, P., Pavlidis, P. & Stamatakis, A. A general species delimitation method with applications to phylogenetic placements. Bioinformatics 29, 2869–2876 (2013).

36. Puillandre, N., Lambert, A., Brouillet, S. & Achaz, G. ABGD, Automatic Barcode Gap Discovery for primary species delimitation. Mol. Ecol. 21, 1864–1877 (2012).

37. Meier, R., Shiyang, K., Vaidya, G. & Ng, P. K. L. DNA barcoding and taxonomy in Diptera: a tale of high intraspecific variability and low identification success. Syst. Biol. 55, 715–728 (2006).

38. Wang, X. P. et al. Phylogenetic relationships among the colobine monkeys revisited: new insights from analyses of complete mt genomes and 44 nuclear non-coding markers. PLoS One 7(4),e36274 (2012).

39. Napier, J. R. & Napier, P. H. A Handbook of Living Primates. Academic Press, London (1967).

40. Tilson, R. L. Infant coloration and taxonomic affinity of the Mentawai Islands leaf monkeys, *Presbytis potenziani*. J. Mammal. 57, 766–769 (1976).

41. Weitzel, V. A preliminary analysis of the dental and cranial morphology of *Presbytis* and *Trachypithecus* in relation to diet. MA thesis, Australian National University, Canberra (1983).

42. Voris, H. K. Maps of Pleistocene sea levels in Southeast Asia: shorelines, river systems and time durations. J Biogeogr 27, 1153–1167 (2000).

43. Slik, J. W. et al. Soils on exposed Sunda Shelf shaped biogeographic patterns in the equatorial forests of Southeast Asia. PNAS 108, 12343–12347 (2011).

44. Roos, C., Nadler, T. & Walter, L. Mitochondrial phylogeny, taxonomy and biogeography of the silvered langur species group (*Trachypithecus cristatus*). Mol. Phylogenet. Evol. 47, 629–636 (2008).

45. Tan, S. H. D., Ali, F., Kutty, S. N. & Meier, R. The need for specifying species concepts: how many species of silvered langurs (*Trachypithecus cristatus* group) should be recognized? Mol. Phylogenet. Evol. 49, 688–689 (2008).

46. Ang, A. Final Report (Phase 1: 2016-2018): Species Action Plan for the Conservation of Raffles’ Banded Langur (*Presbytis femoralis femoralis*) in Malaysia and Singapore. Unpublished Report, Wildlife Reserves Singapore (2018).

47. Shevade, V. S., Potapov, P. V., Harris, N. L. & Loboda, T. V. Expansion of industrial plantations continues to threaten Malayan tiger habitat. Remote Sens. 9(7),747 (2017).

48. Shevade, V. S. & Loboda, T. V. Oil palm plantations in Peninsular Malaysia: Determinants and constraints on expansion. PLoS One 14(2),e0210628 (2019).

49. Rizaldi et al. Preliminary study on the distribution and conservation status of the East Sumatran banded langur (*Presbytis femoralis percura*) in Riau Province, Sumatra, Indonesia. Asian Primates J. 8(1),25–36 (2019).

50. Uryu, Y. et al. Sumatra’s forests, their wildlife and the climate, windows in time: 1985, 1990, 2000, and 2009. WWF-Indonesia Technical Report Jakarta Indonesia. (2010).

51. World Bank. The cost of fire: An economic analysis of Indonesia’s 2015 fire crisis. Indonesia Sustainable Landscapes Knowledge Note. 1(2016).

52. Nijman, V. & Manullang, B. *Presbytis melalophos*. In: The IUCN Red List of Threatened Species 2008: e.T18129A7666452. Accessed on 12 September 2019 (2008).

53. Chiou, K. L. & Bergey, C. M. Methylation-based enrichment facilitates low-cost, noninvasive genomic scale sequencing of populations from feces. Sci. Rep. 8, 1975 (2018).

54. Burgin, C. J., Colella, J. P., Kahn, P. L. & Upham, N. S. How many species of mammals are there? J. Mammal. 99, 1–14 (2018).

55. Nsubuga, A. M. et al. Factors affecting amount of genomic DNA extracted from ape faeces and the identification of an improved sample storage method. Mol. Ecol. 13, 2089–2094 (2004).

56. Hahn, C., Bachman, L. & Chevreux, B. Reconstructing mitochondrial genomes directly from genomic next-generation sequencing reads-a baiting and iterative mapping approach. Nucleic Acids Res. 41, e129 (2013).

57. Langmead, B. & Salzberg, S. L. Fast gapped-read alignment with Bowtie 2. Nat. Methods 9, 357–359 (2012).

58. Li, H. et al. The Sequence Alignment/Map format and SAMtools. Bioinformatics 25, 2078–2079 (2009).

59. Wilm, A. et al. LoFreq: a sequence-quality aware, ultra-sensitive variant caller for uncovering cell-population heterogeneity from high-throughput sequencing datasets. Nucleic Acids Res 40, 11189–11201 (2012).

60. Isokallio, M. A. & Stewart, J. Low-frequency variant calling from high-quality mtDNA sequencing data. protocols.io doi: 10.17504/protocols.io.nfkdbkw (2018).

61. McKenna, A. et al. The Genome Analysis Toolkit: a MapReduce framework for analyzing next-generation DNA sequencing data. Genome Res. 20, 1297–1303 (2010).

62. Bernt, M. et al. MITOS: Improved de novo Metazoan mitochondrial genome annotation. Mol. Phylogenet. Evol. 69, 313–319 (2013).

63. Kumar, S., Stecher, G., Li, M., Knyaz, C. & Tamura, K. MEGA X: molecular evolutionary genetics analysis across computing platforms. Mol. Biol. Evol. 35, 1547–1549 (2018).

64. Katoh, K., Kuma, K., Hiroyuki, T. & Miyata, T. MAFFT version 5: improvement in accuracy of multiple sequence alignment. Nucleic Acids Res. 33, 511–518 (2005).

65. Vaidya, G., Lohman, D. J. & Meier, R. SequenceMatrix: concatenation software for the fast assembly of multi‐gene datasets with character set and codon information. Cladistics 27, 171–180 (2011).

66. Stamatakis, A. RAxML version 8: a tool for phylogenetic analysis and post-analysis of large phylogenies. Bioinformatics 30, 1312–1313 (2014).

67. Lanfear, R., Frandsen, P. B., Wright, A. M., Senfeld, T. & Calcott, B. PartitionFinder 2: new methods for selecting partitioned models of evolution for molecular and morphological phylogenetic analyses. Mol. Biol. Evol. 34, 772–773 (2016).

68. Srivathsan, A. & Meier, R. On the inappropriate use of Kimura‐2‐parameter (K2P) divergences in the DNA‐barcoding literature. Cladistics 28, 190–194 (2012).

69. Bouckaert, R. et al. BEAST 2: A software platform for Bayesian evolutionary analysis. PLoS Comput. Biol. 10(4),e1003537 (2014).

70. Stone, A. C. et al. More reliable estimates of divergence times in Pan using complete mtDNA sequences and accounting for population structure. Phil. Trans. R. Soc. Lond. B Biol. Sci. 365(1556),3277–3288 (2010).

71. Bouckaert, R. R. & Drummond, A. J. bModelTest: Bayesian phylogenetic site model averaging and model comparison. BMC Evol. Biol. 17, 42 (2017).

72. Miller, G. S. Jr. The langurs of the *Presbytis femoralis* group. J. Mammal. 15, 124–137 (1934).

73. Elliot, D. G. Subgenus *Pygathrix*. In: A Review of the Primates, Vol. 3. American Museum of Natural History, New York (1913).

74. Miller, G. S. Jr. Fifty-one new Malayan mammals. Smithsonian Miscellaneous Collections 61, 28 (1913).

75. Pocock, R. I. The monkeys of the genera *Pithecus* (or *Presbytis*) and *Pygathrix* found to the east of the Bay of Bengal. Proc. Zool. Soc. Lond. 104(4),895–961 (1934).

76. Raven, H. C. Wallace’s Line and the Distribution of Indo-Australian Mammals. New York Press (1935).

77. Chasen, F. N. A handlist of Malayan mammals. B. Raffles Mus. 15, 1–209 (1940).

78. Hooijer, D. A. Quaternary langurs and macaques from the Malay Archipelago. Zoologische Verhandelingen Uitgegeven door het Rijksmuseum van Natuurlijke Historie te Leiden 55, 1–64 (1962).

79. Medway, L. The monkeys of Sundaland: ecology and systematics of the cercopithecids of a humid equatorial environment. In: Napier, J. R. & Napier, P. H. (eds). Old World Monkeys: Evolution. Academic Press, New York, 513–553 (1970).

80. Thorington, R. W. Jr. & Groves, C. P. An annotated classification of the Cercopithecoidea (1970). In: Napier, J. R. & Napier, P. H. (eds). Old World Monkeys: Evolution. Academic Press, New York (1970).

81. Wilson, C. C. & Wilson, W. L. Behavioral and morphological variations among primate population in Sumatra. Yearb. Phys. Anthropol. 20, 207–233 (1977).

82. Medway, L. The Wild Mammals of Malaya (Peninsular Malaysia) and Singapore. Oxford University Press, Oxford (1983).

83. Brandon-Jones, D. Colobus and leaf monkeys. In: MacDonald, I. D. (ed). Encyclopaedia of Mammals. George Allen and Unwin, London, 398–408 (1984).

84. Napier, P. H. Catalogue of Primates in the British Museum (Natural History) and elsewhere in the British Isles. Part III: Family Cercopithecidae, Subfamily Colobinae. British Museum (Natural History), London (1985).

85. Aimi, M. & Bakar, A. Taxonomy and distribution of *Presbytis melalophos* group in Sumatra, Indonesia. Primates 33, 191–206 (1992).

86. Brandon-Jones, D. et al. Asian primate classification. Int. J. Primatol. 25, 97–164 (2004).

87. Md.-Zain, B. M. Molecular systematic of the genus *Presbytis*. PhD thesis. Columbia University, New York (2001).

