## Supplementary Figures for "Faecal DNA to the rescue: Shotgun sequencing of non-invasive samples reveals two subspecies of Southeast Asian primates to be Critically Endangered species"

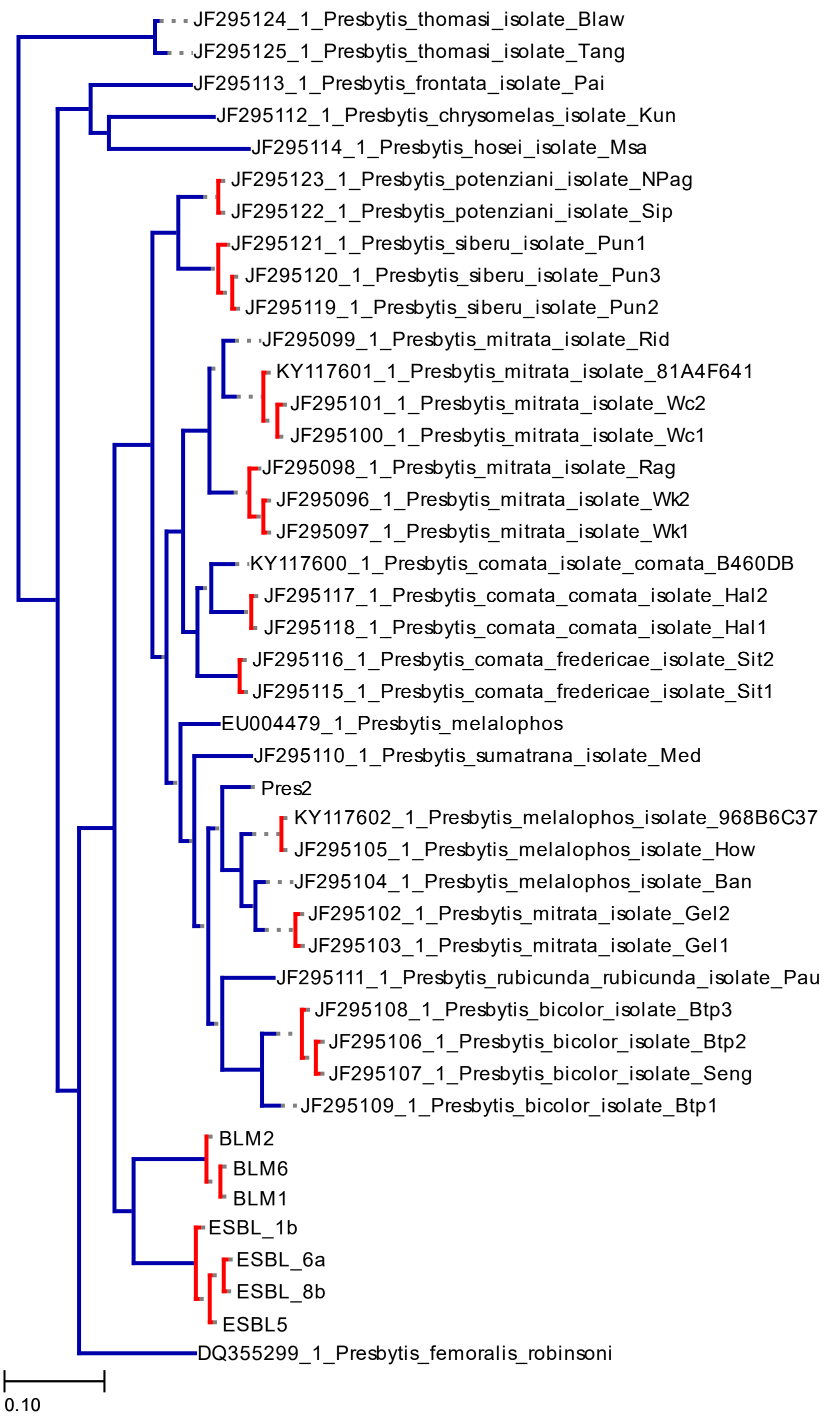


Figure S1: Species delimitation using PTP based on *Presbytis* mitogenome+Cytb+ HV1 dataset

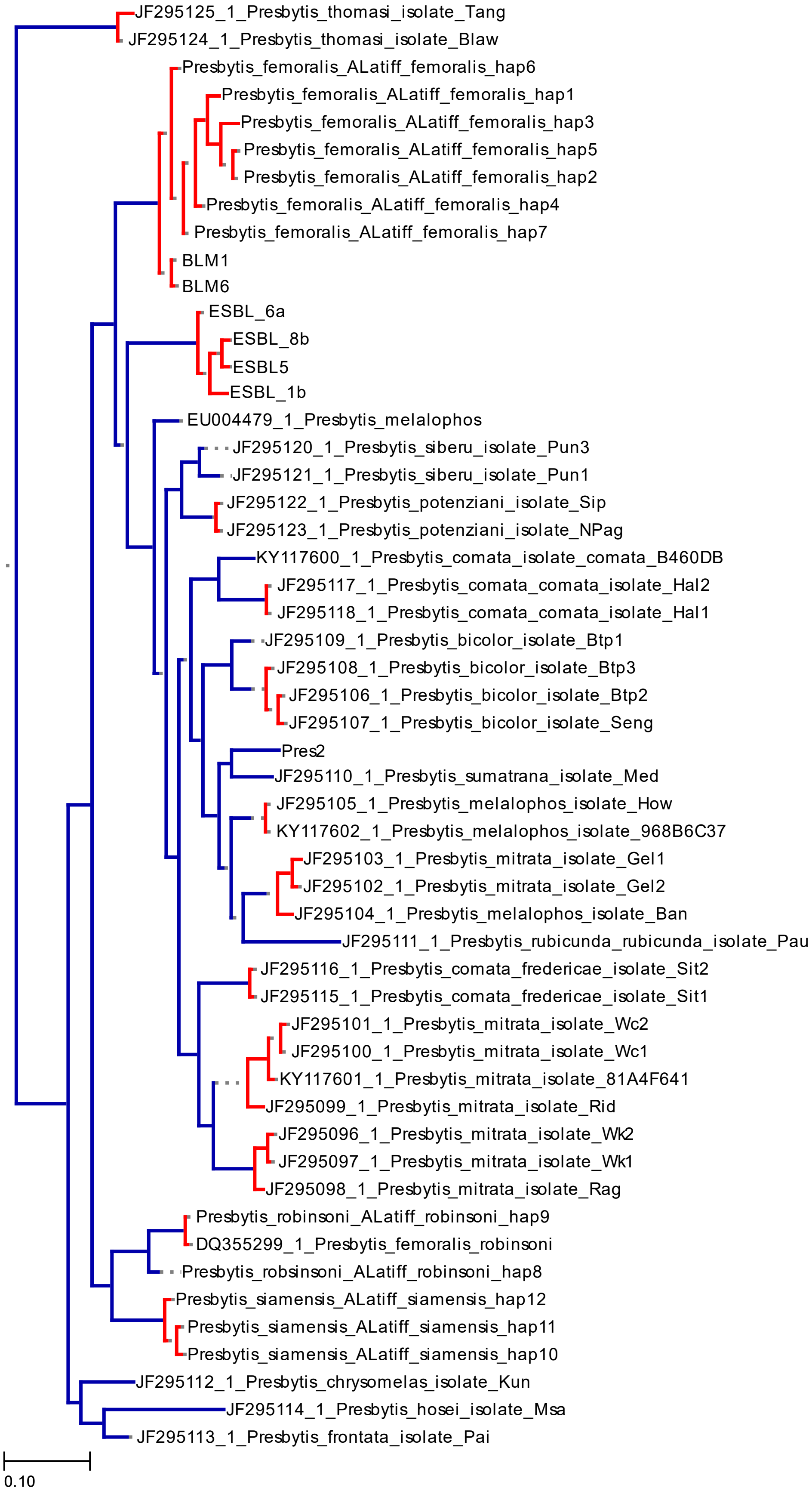


Figure S2: Species delimitation using PTP based on *Presbytis* HV1 only dataset

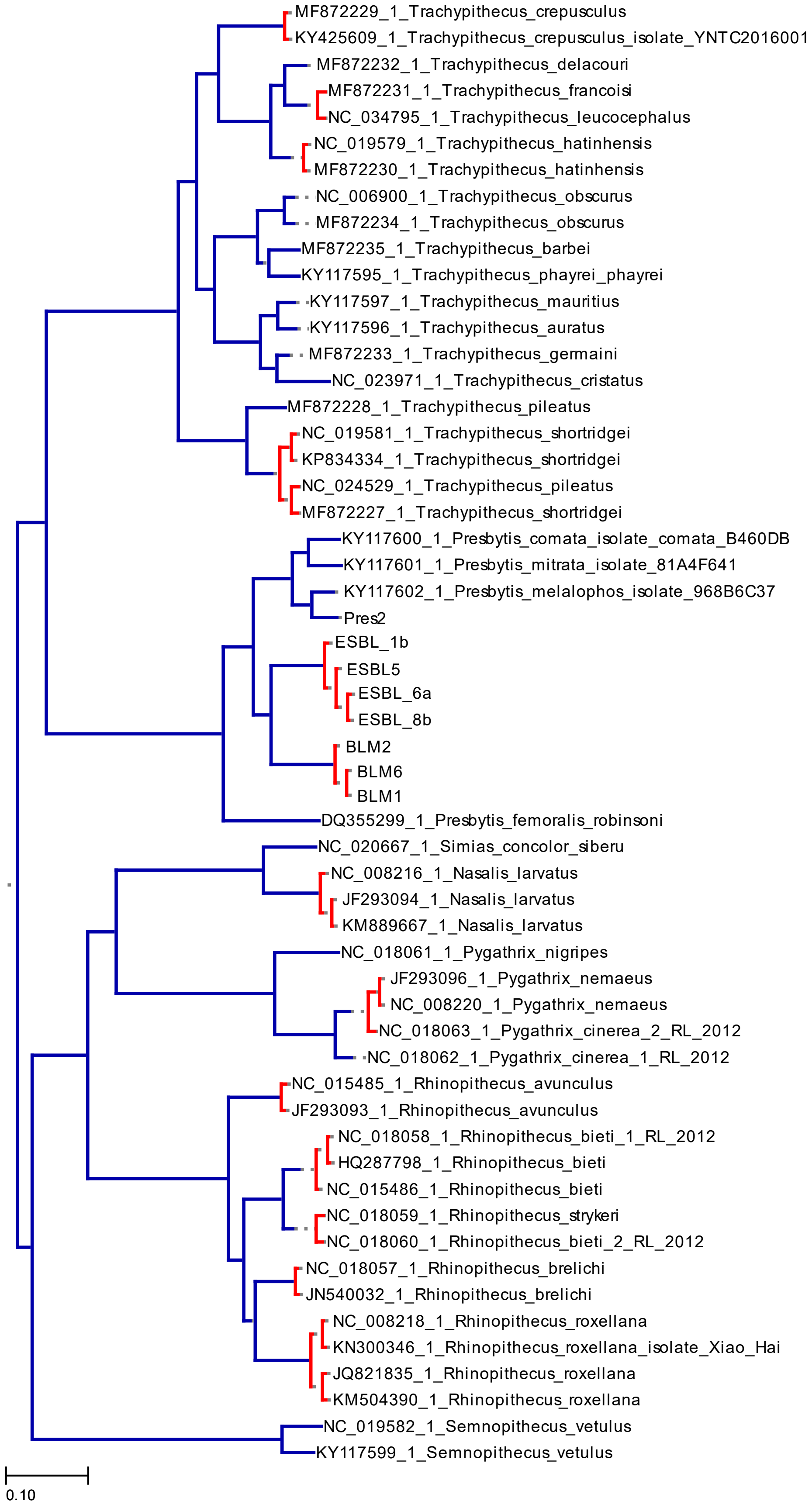


Figure S3: Species delimitation using PTP based on Asian colobine mitogenome dataset

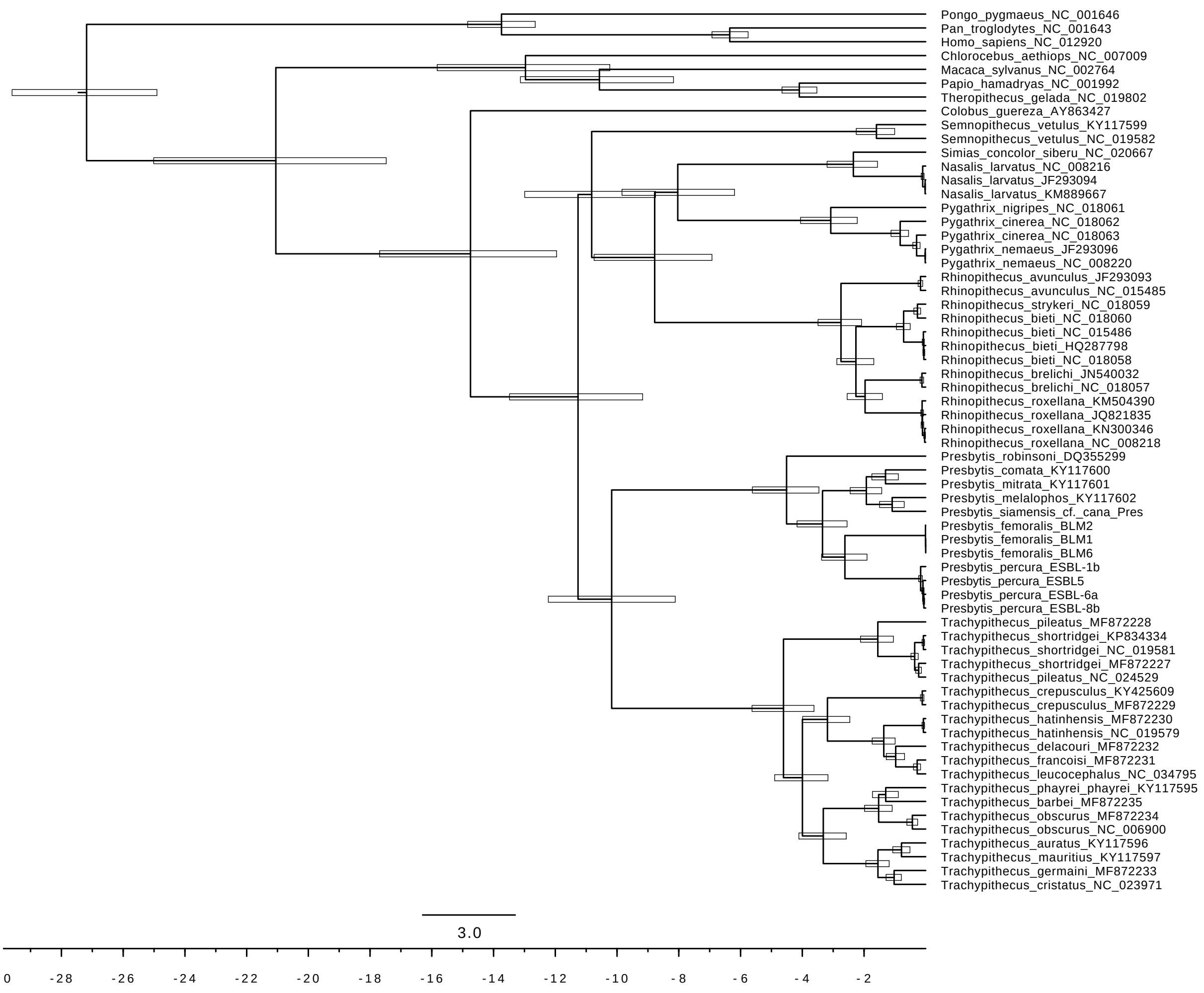


Figure S4. Divergence times estimates based on mitochondrial genomes for Asian colobine primates (partitioned by gene)

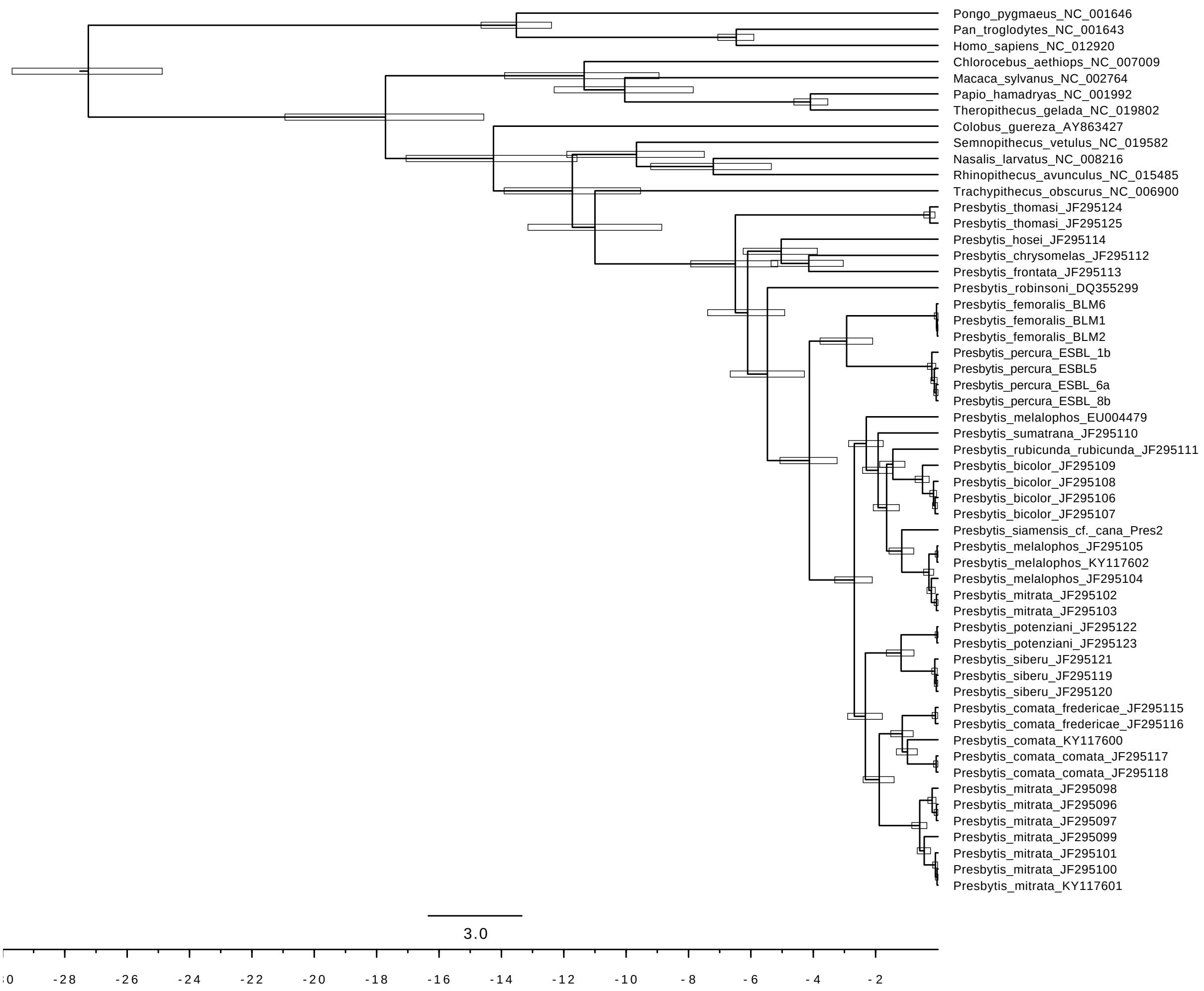


Figure S5 Divergence times estimates based on cyt-b for *Presbytis*.
